# Genome evolution in a putatively asexual wasp

**DOI:** 10.1101/2020.12.23.424202

**Authors:** Eric S. Tvedte, Austin C. Ward, Benjamin Trendle, Andrew A. Forbes, John M. Logsdon

## Abstract

Asexual lineages are destined for extinction—a result predicted by theory and revealed in practice. Short-term benefits of asexuality are eventually outstripped by their fitness costs: losses of sex and recombination are together expected to reduce efficacy of selection, increase mutation load, and thus, lower fitness. We characterized genomic patterns of accumulating mutations in *Diachasma muliebre*, a parasitic wasp that has apparently lost sex, an inference supported by many decades of field collections of 1000s of individuals in which only females were found. The split between *D. muliebre* and its closest sexual relative, *Diachasma ferrugineum*, is quite recent, allowing us to observe initial events in the evolution of this putative asexual species. First, we find a faster rate of molecular evolution across the *D. muliebre* genome. In addition, we observed a marked excess of replacement nucleotide substitutions in orthologous genes in the putatively asexual *D. muliebre* when compared to *D. ferrugineum*. This pattern directly indicates genome-wide relaxed selection in this young, putatively asexual species, the resulting mutational load from which is expected to ultimately lead to extinction. However, these genomic effects occur in the presence of genomic recombination initially detected by a previous study and also supported by analyses of genome-wide substitution rates within codons. In addition, following completion of the genome sequence and its analysis, we discovered two *D. muliebre* males, suggesting the possibility of rare sex in this species. Haplodiploid animals, including the sexual ancestors of *D. muliebre*, bear small genetic loads, likely making their initial transitions to asexuality relatively benign. Paradoxically, an elevated rate of mutation accumulation resulting from asexuality, when accompanied by retention of recombination and/or rare sex, could actually be beneficial: we hypothesize that the novel variation introduced by mutation along with limited shuffling of genes may facilitate initial adaptation and extend persistence of such lineages.

## Introduction

Sexual reproduction is omnipresent among eukaryotes despite its considerable metabolic and genetic costs. This apparent paradox has generated substantial discourse regarding the presumed benefits of sex (Otto 2009; Hartfield and Keightley 2012). The ubiquity of sex and the “twiggy” representation of asexual lineages on phylogenetic trees have been interpreted as signals that transitions to asexuality are always accompanied by major fitness costs—and that asexuality is eventually an evolutionary dead end (Maynard Smith 1978; Bell 1982).

Explanations of why fitness would decrease after a transition to asexuality usually invoke the increase in genetic load of deleterious mutations in asexual genomes (Lynch et al. 1993). Without outcrossing and meiotic recombination, asexuals are predicted to experience limitations on the efficacy of selection to act independently within and among loci and to expunge deleterious alleles (Hill and Robertson 1966; Felsenstein 1974). Moreover, asexual individuals with low genetic loads may often be lost *via* drift and the inability to restore these least loaded genotypes *via* recombination should lead to the irreversible accumulation of deleterious mutations within an asexual lineage (Muller 1964; Charlesworth and Charlesworth 1997). Overall, high genetic loads experienced by asexual lineages is expected to limit their ability to persist over long evolutionary timespans.

The absence of meiotic recombination and the resulting reductions in efficacy of selection in asexuals is predicted to produce a distinct genomic signal: a genome-wide, increased ratio of nonsynonymous to synonymous differences between species (Glémin and Galtier 2012). Previous studies provide some support for a faster accumulation of putatively deleterious mutations in asexuals; however, until recently most have examined just a small number of genes, often rendering the results equivocal (Hartfield 2016). Recent transcriptome and whole-genome analyses have improved the amount of data bearing on this question, but have also shown mixed support for increased mutation accumulation in asexual lineages (Jaron et al. 2020). Some studies have suggested a reduced efficacy of purifying selection to remove harmful mutations in clonal asexual lineages (Hollister et al. 2014; Lovell et al. 2017; Bast et al. 2018), while others show no effect (Ament-Velásquez et al. 2016; Lindsey et al. 2018; Brandt et al. 2019) or even more effective selection in asexuals (Brandt et al. 2017). [It should be noted that we use “mutation accumulation” to describe differences between species in this context instead the cumbersome, but more accurate, “substitution accumulation”.]

Why have previous studies not detected a consistent mutational signal in asexual genomes, if it exists? Comparisons between sexual and asexual sister species are difficult due to uncertainties about the age of the split (Ament-Velásquez et al. 2016; Brandt et al. 2019) (and, for comparisons among asexual groups, variability in asexual age), or they are complicated by hybridization and/or polyploidy in the transition event (Hollister et al. 2014; Ament-Velásquez et al. 2016). It is also possible that some presumed asexuals are, in fact, not entirely asexual, but instead are capable of sexual reproduction, albeit occasionally, or even rarely (Schurko et al. 2009). Nonetheless, a number of genomes from presumed asexuals have been sequenced and compared to their sexual relatives, as summarized recently (Jaron et al. 2020). Notably, no consistent feature—including mutation accumulation—is shared among all (or even a majority) of the 24 presumptive asexual animal genomes that Jaron *et al*. (2020) surveyed. Some other unusual features found among some asexual animal genomes include changes in heterozygosity, increased numbers of horizontally transferred genes, reductions in transposable element content, and loss of genes involved in sexual reproduction. Although these data may indicate no feature shared in common (and diagnostic for) asexual genomes studied to date, many of these odd aspects are predicted to occur only following the long-term persistence of asexuality. Indeed, it is possible that comparisons among asexual genomes with different divergence times from their sexual ancestors obscures potential genomic commonalities, in particular those that might arise either in late or early stages of asexuality.

An ideal system to assess the initial impacts of asexuality on a genome would be one in which the transition to asexuality is recent, where the ecology is little changed, and for which genomic data are readily available for evolutionary study (Tvedte et al. 2019a). The presumed asexual parasitic wasp *Diachasma muliebre* split from its sexual relative *Diachasma ferrugineum* in an apparent single loss-of-sex event 10,000YA-1MYA (Forbes et al. 2013)(**Supplemental Information**), but the two wasps otherwise differ very little, attacking sister species of specialist tephritid fruit flies on cherry trees. Although details of the *D. muliebre* reproductive life cycle are essentially unknown, separate lines of evidence are consistent with it being asexual, including: a) in >2300 individual specimens from ~70 years of field collections and across its entire geographic range, only females have been collected and not a single male was identified prior to this work (see **Results & Discussion** and **Supplemental Information**; **Table S1**)—though field collections analyzed in 2020 revealed the presence of two apparent *D. muliebre* males and b) there appears to be complete association between mitochondrial and nuclear genotypes (Forbes et al. 2013).

The loss of sexual reproduction has both short- and long-term consequences which may be revealed in the genomes of asexual lineages. Here, our comparative genomic analyses of the putative asexual wasp *D. muliebre* and its close sexual relative, *D. ferrugineum* show striking patterns of sequence divergence associated with a recent transition to asexuality and consistent with current evolutionary theory. Notably, these patterns emerge and persist even in the presence of detectable genomic recombination and the presence of rare males in this species.

## Results & Discussion

We obtained Illumina sequencing datasets from the genomes of both *D. muliebre* and *D. ferrugineum* (**Table S2**) and compared these with existing genome sequences from their next closest sexual relative (*Dichasma alloeum*) and two other sexual braconid wasps (*Fopius arisanus, Microplitis demolitor*). After considerable annotation and filtering (see **Materials & Methods**), we recovered 3,127 genes with full coding DNA sequences (CDS) coverage in all five wasp species, representing ~25% of the predicted genes in the *D. alloeum* genome (Tvedte et al. 2019b).

In the absence (or decreased functionality) of sex and canonical recombination in asexuals, the proportion of mutation-free genes is predicted to decrease and, by inference, their mutational load will increase. Consistent with this prediction, we find that gene sequences from *D. muliebre* are more different than genes from *D. ferrugineum* when both are compared to *D. alloeum*. Indeed, 1,219 of the 3,127 genes have higher pairwise distances in *D. muliebre* (pdistDM > pdistDF), while 899 genes have higher pairwise distances in *D. ferrugineum* (pdistDM < pdistDF) (**Figure 1**; *χ^2^* = 48.35, p < 0.0001). Additionally, 149 genes are identical between *D. ferrugineum* and *D. alloeum* (pdistDF = 0), compared to 133 identical genes between *D. muliebre* and *D. alloeum* (pdistDM = 0).

**Figure 1.**
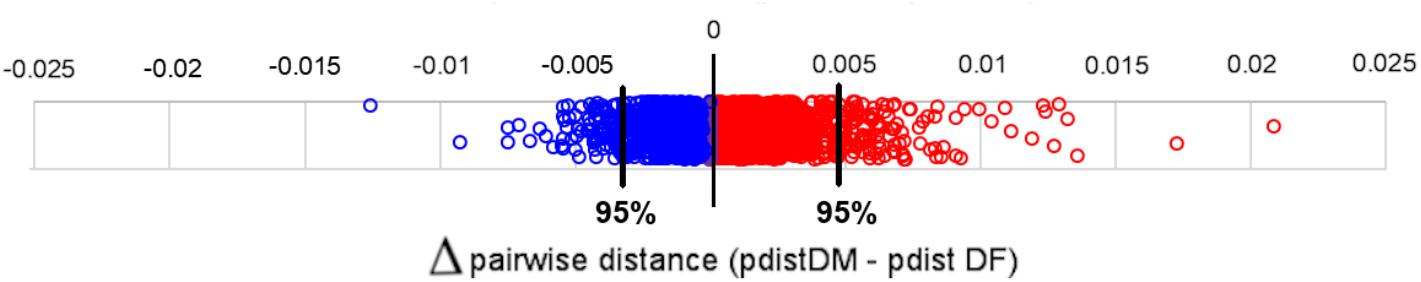
Relative pairwise distances in sexual and asexual *Diachasma*. Positive (red) values indicate genes with more nucleotide differences in asexuals (pdistDM > pdistDF), negative (blue) values indicate genes with more differences in sexuals (pdistDM < pdistDF), zero (purple) values indicate genes with equivalent distances (pdistDM = pdistDF). Bolded black lines correspond to the 90th percentile of positive and negative Δ values. All pairwise distance values were calculated against a common wasp reference (*D. alloeum*).

Despite the recent origin of this asexual wasp, comparisons of pairwise distances across the entire gene set display a prominent shift of the *D. muliebre* mutation landscape relative to *D. ferrugineum*, reflecting a faster rate (accumulation) of change in the asexual (**Figure 2, Figure S1**). There is a significant difference between mean pairwise distances for *D. muliebre* genes (MpdistDM = 0.005496, SD = 9.365E-6) relative to *D. ferrugineum* genes (MpdistDF = 0.005089, SD = 7.953E-6) when compared to *D. alloeum* homologs; this indicates more new mutations persist in the asexual lineage (Wilcoxon rank sum p = 4.17E-7). Although the distinct mutational landscapes in sexual and asexual wasps could be confounded by the non-independence of genes contained in linkage groups, the sampled scaffolds did not show evidence of substantial linkage among genes in our dataset (**Supplemental Information**).

**Figure 2.**
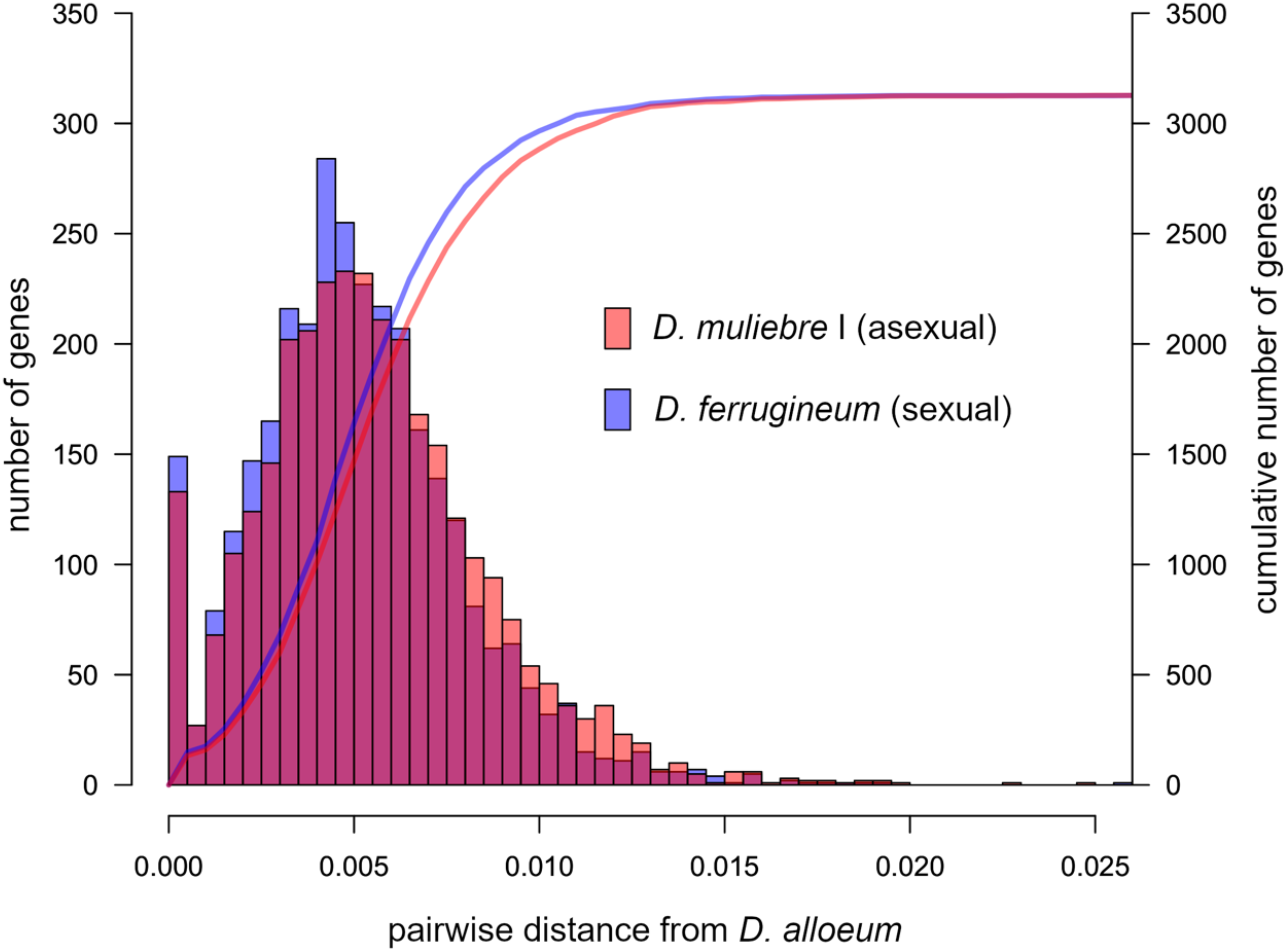
Mutation landscapes of sexual and asexual *Diachasma*. Columns represent binned frequencies (left y-axis) of corresponding p-distance values from *D. alloeum*. Curves represent the cumulative frequency of binned columns (right y-axis).

To further evaluate patterns of mutation accumulation in *Diachasma* genomes, we combined coding sequences from all 3,127 genes into a concatenated dataset. Given the recency of the presumed loss-of-sex event in these wasps, the signal generated by differences in rates of mutation accumulation is likely to still be small, and therefore comparisons involving single or few genes may not be informative (Tvedte et al. 2017). A maximum likelihood analysis of the concatenated coding sequences displays a clear increase in the overall rate of molecular evolution in *D. muliebre* relative to *D. ferrugineum* (**Figure 3A**). Indeed, the total number of unique differences in *D. muliebre* is greater than in *D. ferrugineum* (χ^2^ = 256.91, p < 0.0001) consistent with observations from pairwise distance data (**Table 1**).

**Figure 3.**
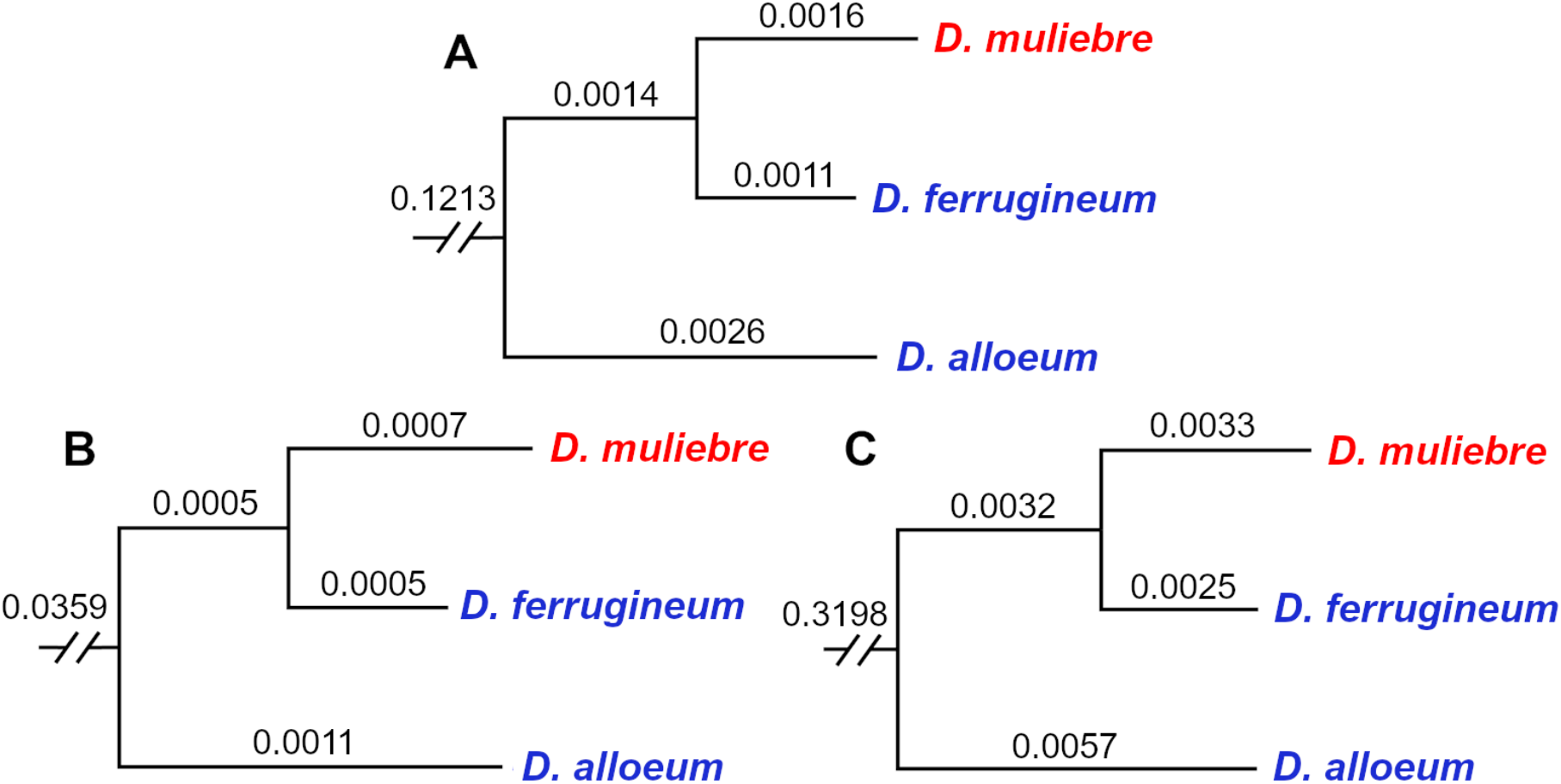
Maximum likelihood analysis of concatenated nuclear gene dataset in *Diachasma*. Asexual (red) and sexual (blue) *Diachasma* lineages are shown, and informative branch lengths are labeled. **A.** Phylogenetic tree of all sites (3,715,368 bp). **B.** Phylogenetic tree of 1^st^/2^nd^ codon sites (2,476,912 bp). **C.** Phylogenetic tree of 3^rd^ codon sites (1,238,456 bp).

**Table 1.** Relative rate analysis of concatenated gene set in *Diachasma*.

| #<br>genes | length<br>(bp) | number of unique differences |  |  |  |  |  |
| --- | --- | --- | --- | --- | --- | --- | --- |
|  |  | all sites |  | 1<sup>st</sup> / 2<sup>nd</sup> codon<br>positions |  | 3<sup>rd</sup> codon positions |  |
|  |  | DF | DM | DF | DM | DF | DM |
| 3,127 | 3,715,368 | 4,148 | **5,742<sup>a</sup>** | 1,113<br>(26.83%) | **1,702<sup>a</sup>**<br>(29.64%) | 3,035<br>(73.17%) | **4,040<sup>a</sup>**<br>(70.36%) |

*D. alloeum* was used as the outgroup to measure the number of unique differences in the sexual *D. ferrugineum* (DF) vs. asexual *D. muliebre* (DM) lineages. 14,598 sites were shared by both DM and DF and 36 sites were unique in all three species. <sup>a</sup> $\chi^2$ test statistic was statistically significant ( $p \ll 0.005$ ), *i.e.* the bolded value represents a significantly higher number of unique differences than expected under a null model of equal evolutionary rates.

To distinguish between effects generated by accelerated mutation rate *per se* in the asexual and reduced efficacy of selection, we also evaluated these combined datasets by codon position. An increase in the intrinsic mutation rate in an asexual lineage would be characterized by a greater number of mutational changes at selectively neutral sites (*e.g*. synonymous sites). We analyzed the aligned 3^rd^ codon positions to estimate this effect. Similarly, if asexuality is associated with the reduction in the efficacy of selection to act independently on physically proximate loci to remove mildly deleterious mutations, we expect to observe the increased incidence of substitutions at nonsynonymous sites, estimated here using 1^st^ and 2^nd^ codon positions. As a result of this increased rate of mutation in the asexual wasps, *D. muliebre* shows a longer branch length relative to *D. ferrugineum* when using 3^rd^ codon position partitions, and a relative rate test indicated a significant excess of unique differences in *D. muliebre* (χ^2^ = 142.76, p < 0.00001) (**Table 1, Figure 3B**). Similar to 3^rd^ codon position observations, *D. muliebre* had longer branch lengths compared to those of its closest sexual relative, *D. ferrugineum* (**Figure 3C**). The number of unique differences was substantially higher in *D. muliebre* at 1^st^/2^nd^ positions (χ^2^ =123.24, p < 0.00001), and the asexual genome also contained a larger proportion of total unique differences occurring at 1^st^/2^nd^ codon positions (**Table 1**). We tested for selective pressures on synonymous sites by measuring the effective number of codons (ENC) and codon deviation coefficients (CDC) and our results suggest that accelerated evolution in *D. muliebre* is not explained by relaxed selection on codon usage (**Table S3, Supplemental Information**). Taken together, this pattern suggests that *D. muliebre* possesses a larger mutational load relative to *D. ferrugineum*, with putatively deleterious mutations more often retained in the asexual lineage.

To further consider the possible effects of asexuality on individual genes, we extended our analyses to compare the rates of molecular evolution between *D. muliebre* and *D. ferrugineum* within and among each of the 3,127 genes. Our comparisons of the amino-acid replacement (nonsynonymous) rates of substitution (dN) between genes revealed results that are mutually complementary. First, there are significantly more genes in the zero class of estimated replacement substitutions in the sexual species (*D. ferrugineum*) compared to the asexual (**Figure 4**). Second, there are significantly more genes having replacement substitutions (χ^2^ = 5.303, p = 0.0203) in the asexual (*D. muliebre*) compared to the sexual species. Third, there are significantly more replacement substitutions (estimated by 1^st^ and 2^nd^ position differences) in *D. muliebre* (χ^2^ = 33.006, p < 0.0001), even when correcting for the fact that there are more overall differences in *D. muliebre* (χ^2^ = 4.476, p = 0.0344). Strikingly, the pattern in **Figure 4** shows a clear shift, further indicating reduction in the efficacy of selection in the asexual. Our parallel analyses of synonymous (dS), did not reveal a significant pattern of differences between the sexual and asexual, beyond an overall excess seen in *D. muliebre*—which is expected given that *D. muliebre* has more overall substitutions (**Figure S2, Supplemental Information**). These results provide a clear snapshot of the initial processes of molecular evolution near the onset of asexuality: genes that are evolving under varying degrees of purifying selection—from the strongest to the weakest—show an increased rate of replacement substitution in the asexual. This effect appears to be general and genome-wide; however, further analyses will be needed to assess this among and between genes in various functional classes.

**Figure 4.**
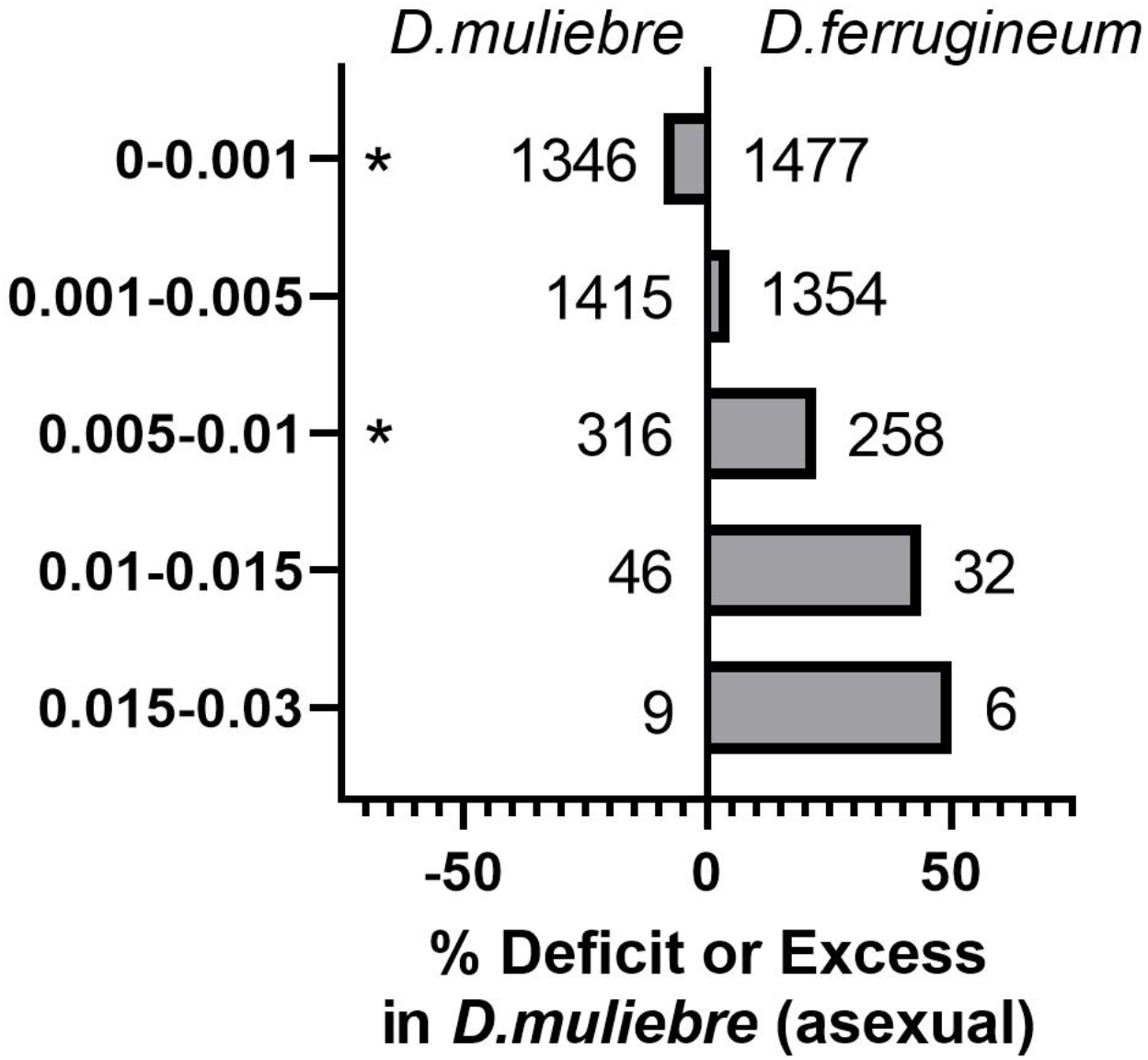
Comparison of replacement substitutions (dN) in 3,127 orthologs from *D. muliebre* and *D. ferrigineum*, compared to *D. alloeum*. Replacement substitutions (dN) were calculated in PAML comparing *D. muliebre* and *D. ferrigineum* to *D. alloeum* in three species alignments. The total number of genes in each bin for each species is shown; for example, there are 1346 and 1477 genes with dN = 0-0.001 (all with identical amino acid sequences observed) in *D.m*. and *D.f.*, respectively. For each bin, the percent excess (or deficit) in *D. muliebre* is shown as a percentage compared to the number of genes for each class observed in *D. muliebre*. The bins marked with asterisks (*) are significantly different (see text).

In addition to the overall increase in the rate of molecular evolution resulting from reduced selection in an asexual, both the efficacy and pattern of selection will be drastically altered across the genome. In fact, for asexuals that are not experiencing recombination, their entire genome is inherited clonally, disallowing mutations in adjacent sites in the genome to be selected independently—even if their selective effects differ. Thus, in asexuals, substitutions at silent *versus* non-silent codon positions would not directly follow the familiar patterns observed in sexual species. Given that rates of new mutations at 1^st^, 2^nd^ and 3^rd^ positions in codons are approximately equal within a species, their rates of substitution after the onset of clonality would only reflect the removal of deleterious mutations along with all linked mutations (on that chromosome). Then, the prediction in a clonal asexual will be an increase in the observed ratio of non-silent: silent codon substitutions across the genome relative to the ancestral sexual state. However, since recombination has previously been inferred in *D. muliebre* (Forbes et al. 2013; Tvedte et al. 2017)—a result which may be related to the presence of rare males (see below)— the prediction is that substitution patterns within codons in this asexual should be more similar to patterns in sexuals than to a true clonal asexual. In fact, the ratio of observed substitutions at 1^st^ + 2^nd^ positions *versus* 3^rd^ in *D. muliebre* is 0.42, compared to 0.37 in *D. ferrugineum* (**Table 1**). These data further indicate that *D. muliebre* is experiencing a measurable decrease in the genome-wide efficacy of purifying selection (on the order of ~15%), discounting possible effects of positive selection (which would be predicted, on average, to be similar in both species). When comparing the inferred synonymous versus non-synonymous rates across all 3,127 genes, the mean dN minus mean dS is 20% greater in *D. muliebre* (data not shown), consistent with the observed patterns by codon position. Notably, the similarity of this ratio of observed substitutions between the sexual and asexual wasps is consistent with the presence of recombination in *D. muliebre* (where the extreme predicted ratio in a clonal asexual with no recombination should approach 2.0); however, the generality of this result is unknown, as we are unaware of other such analyses in the literature. Two major hypotheses could explain why substitution rates within codons vary somewhat independently, albeit less so than for a sexual: 1) Perhaps *D. muliebre* is not, in fact, strictly asexual. The recent discovery of rare males may occasionally provide opportunity for infrequent sex and meiotic recombination; 2) Sufficient meiotic recombination could be provided by automictic reproduction in *D. muliebre*. Prior to the recent discovery of males, we favored the second hypothesis given the data currently available (Forbes et al. 2013; Tvedte et al. 2017); however, this will require further investigation, especially focusing on the biological relevance of *D. muliebre* males (see below and **Supplemental Information**).

In any case, the clear result of increased rate of genome-wide substitution patterns we observe in *D. muliebre* are themselves strong predictors for asexuality and are not consistent with sex. Nonetheless, the recent discovery of two *D. muliebre* males in 2020 field collections has required us to consider possible implications for asexuality in this species. First, given the extreme rarity of finding males in this species (only 2 males in over 2800 wasps collected over ~70 years), it is likely that the vast majority of reproduction in this species is asexual. It is possible that these are an example of “spanandric”, non-functional, males that are rarely produced in some otherwise obligately asexual species (Smith et al. 2006). Alternatively, these males could be functional and only rarely participate in sexual reproduction in this species. Although new questions are raised (see **Supplemental Information**) and additional analyses will be required to definitively address the possibilities, reproduction in *D. muliebre* is primarily, if not entirely, asexual.

In any case, to our knowledge, this study represents the first demonstration of accelerated mutation accumulation in a putatively asexual organism that is apparently capable of some genome-wide recombination. In a recent analysis comparing sexual and automictic asexual *Trichogramma pretiosum* wasps, Lindsey *et al*. (2018) did not find increased rates of protein evolution in asexuals; however, the absence of a sexual outgroup to *T. pretiosum* prohibited genome-wide rate tests at the nucleotide level. Our comparisons across reproductive modes are largely based on single individual wasps as proxies for entire lineages. Our whole-genome NGS datasets from five previously-identified lineages of *D. muliebre* wasps exhibited very low intraspecific variation (**Table S4, Figure S3, Supplemental Information**), such that our conclusions about one asexual individual reasonably extend to the asexual species. However, increased sampling from *D. muliebre* and *D. ferrugineum* wasps—for example to determine ratios of nonsynonymous to synonymous changes that are fixed *vs*. polymorphic—will be needed to draw further conclusions about genomic consequences of asexuality. Both of these ratios were higher in asexual lineages of *Timema* stick insects, demonstrating that putatively deleterious variants are fixed faster and remain polymorphic for longer in asexuals (Bast et al. 2018).

In addition to providing evidence for mutation accumulation in the early stages of asexual evolution, a new proposition emerges from these data: Asexual animals derived from haplodiploid ancestors—already proposed to be initially more successful (Maynard Smith 1978)—in the presence of automixis (and/or rare sex), may also be less vulnerable to the genomic degradation that general theory suggests should plague other asexuals. Haplodiploid arthropods, which are particularly rich in reproductive transitions to obligate asexuality (van der Kooi et al. 2017), possess a wide variety of automictic mechanisms (Suomalainen et al. 1987; Stenberg and Saura 2009). While transitions to asexual reproduction from diploid sexuals might be generally disadvantageous due to frequent uncovering of deleterious recessive alleles, this component of genetic load is likely to initially be much lower in haplodiploids *(e.g. Diachasma)*, where recessive harmful variants are fully expressed in males (Archetti 2004). Indeed, a recent *in silico* study suggested that automictic asexual lineages could possess both greater genetic diversity and lower genetic loads relative to sexual lineages (Engelstädter 2017).

Now consider an emerging asexual lineage with high diversity, low genetic load, and maintenance of meiotic recombination (*via* automixis or rare sex): its risk of genome degradation may be outweighed by its promise for adaptation. Classical hypotheses for the evolutionary maintenance of sex invoke recombination’s activity to bring together advantageous combinations of alleles (Fisher 1930; Muller 1932) or to generate genetic diversity to increase occupancy of heterogeneous niches (Bell 1982). Both of these advantages—typically reserved for sexual organisms—may apply to *D. muliebre*, a haplodipoid asexual that maintains recombination and has a measurable increase in molecular evolutionary rate as a source of new variation. Indeed, while sexual *D. alloeum and D. ferrugineum* exhibit relatively wide phenotypic variation for adult emergence timing (Forbes et al. 2009) (a trait critical to wasp fitness; **Supplemental Information**), individual lineages of *D. muliebre* have apportioned the temporal niche of their sexual relatives: each has shorter, and incompletely overlapping, emergence windows, with some lineages emerging relatively earlier and others relatively later (Forbes et al. 2013). Earlier-emerging lineages of *D. muliebre* have also recently expanded into a new habitat niche following environmental change (Forbes et al. 2013) (**Supplemental Information**). Access to different combinations of genetic and phenotypic variation may thus reduce competition among asexual lineages, equip an asexual to navigate new environments, and allow asexuals to persist for longer than theory has previously suggested.

## Acknowledgements

We thank Amanda Nelson and Wee Yee for help with wasp collections, Gery Hehman for assistance with Illumina library preparation, and Nick Stewart and Laura Bankers for preliminary sequencing efforts of *Diachasma* wasps that motivated the study. We thank Maurine Neiman and Bryant McAllister and anonymous reviewers of a previous version for their insightful comments on the manuscript. This work was funded by a University of Iowa Internal Funding Initiative grant to A.A.F. and J.M.L. and an NSF DEB (1145355) to AAF.

## Author contributions

E.S.T., A.A.F., and J.M.L. conceptualized the study, designed sequencing experiments, and wrote the manuscript. E.S.T and A.C.W. wrote custom scripts for analysis of sequencing data, E.S.T. and B.T. performed analyses on sequencing data.

## Data availability

Illumina sequencing datasets will be made available on the NCBI SRA database (**Table S2**).

## Materials and Methods

### Preparation of *Diachasma* genomic libraries

We obtained individual female and male *Diachasma* specimens for Illumina sequencing extractions. All collections were made by first collecting *Rhagoletis* fly-infested fruits. Fly larvae were allowed to emerge from fruits and pupate in the lab and were then artificially overwintered in a refrigerator. Wasps emerged from some pupae the following year. We collected two *D. ferrugineum* wasps (one female, one male) in Iowa City, IA, U.S.A. (41.39° N, 91.31° W, 204 m elevation), one *D. ferrugineum* male in South Bend, IN, U.S.A. (41.39° N, 91.31° W, 204 m elevation) and five *D. muliebre* females in Roslyn, WA, U.S.A. (47.22° N, 120.99° W, 685 m elevation). The *D. muliebre* individuals were obtained from a larger collection of wasps and represented five different previously-identified mitochondrial cytochrome C oxidase I (COX1) haplotypes (21). We generated RNA libraries for the Iowa City samples using the TruSeq Stranded mRNA kit (Illumina Inc., San Diego, CA) and DNA libraries for the remaining samples using the KAPA Hyper Prep kit (KAPA Biosystems, Wilmington, MA). We sequenced paired-end DNASeq and RNASeq reads (2 x 300) at the University of Iowa using the Illumina MiSeq platform (Illumina Inc., San Diego, CA). After inspecting raw read data, we performed quality trimming on reads using Trimmomatic v.0.32 (Bolger et al. 2014) and visually confirmed trimmed datasets using FASTQC (Babraham Bioinformatics; https://www.bioinformatics.babraham.ac.uk/projects/fastqc/).

### Reference guided genome assembly and variant calling

Given the haplodiploid genetic system of *Diachasma*, we conducted SNP calling pipelines in the nuclear genome with all female samples (one *D. ferrugineum*, five *D. muliebre*). An overview of the Illumina dataset processing pipeline is provided in **Figure S4**. We adopted the GATK’s Best Practices (Van der Auwera et al. 2013) for generating distinct pipelines for the processing of DNASeq *vs*. RNASeq datasets (https://software.broadinstitute.org/gatk/best-practices/; accessed 2017).

There are three major differences in the RNA *vs*. DNA pipeline. First, we used different programs to map trimmed reads to the *D. alloeum* reference genome. We mapped reads passing quality filters to the *D. alloeum* genome assembly Dall1.0 (Tvedte et al. 2019b) (NCBI Accession GCA_001412515.1). We mapped DNASeq reads using Bowtie2 v.2.3.0 (Langmead and Salzberg 2012) and RNASeq reads using TopHat2 v.2.1.1 (Kim et al. 2013). To map reads to the *D. alloeum* genome, we largely used the default parameters for these tools. For TopHat2, we set the mean inner distance between mate pairs to 450 bp (SD of 20 bp), and for Bowtie2, we set the maximum fragment length for valid paired-end alignments to 800, as these values were consistent with the Illumina library insert sizes. Using the ‘genomecov’ and ‘maskfasta’ functions from BEDTools v.2.26.0 (Quinlan and Hall 2010), we generated new *D. alloeum* reference sequences with masked regions that had <2X coverage. Reads were then re-mapped to the masked reference using Bowtie2 and TopHat2.

Second, for the RNASeq dataset, we executed the GATK program SplitNCigarReads, prior to variant calling using HaplotypeCaller. This tool splits mapped reads containing Ns corresponding to splice junctions and is appropriate only for RNA datasets.

Third, we used specific parameters to filter DNASeq and RNASeq variants using GATK VariantFiltration. Assemblers use different systems for defining the mapping quality (MAPQ) value of reads, and manual inspection of alignments indicated a single MAPQ filter value would not work well across variant callsets. We excluded reads with MQ <2 aligned with Bowtie2 and excluded reads with MQ <40 aligned with TopHat2 and BWA-MEM. All other filtering parameters were identical for all datasets, namely: QD >2.0, FS >60.0, MQRankSum <−12.5, ReadPosRankSum <−8.0, DP >4). After mapping, we manually inspected a subset of genes to assess SNP calling accuracy (**Table S5**, **Table S6**, **Supplemental Information**).

### Ortholog identification and CDS dataset generation

We downloaded protein datasets from the *D. alloeum* (Tvedte et al. 2019b), *F. arisanus* (Geib et al. 2017), and *M. demolitor* (Burke et al. 2018) genome assemblies. Using a custom script, we produced nonredundant protein datasets for each species. Protein datasets were submitted to OrthoVenn v.1 (Wang et al. 2015) for ortholog identification. Using 7,910 single-copy gene clusters assigned by OrthoVenn, we retained 3,127 with full CDS coverage in all wasp species compared in this study, including individual *D. ferrugineum* and *D. muliebre* females. Using the GFF file associated with each genome assembly, we ran a custom script to retrieve genome coordinates for all CDS regions. For *D. alloeum* and *F. arisanus*, we provided CDS coordinates as the −L parameter to retrieve CDS for each gene using GATK FastaReferenceMaker. For *D. ferrugineum* and *D. muliebre*, we executed a similar strategy, using the appropriate masked genome as the −R parameter and the VCF file from the GATK Best Practices pipeline as the −V parameter using GATK FastaAlternateReferenceMaker to obtain CDS for these wasp species. For each of the genes in the dataset, we tested whether the asexual lineage *D. muliebre* possesses a greater mutational load relative to the sexual lineage, *D. ferrugineum*. To do this, we produced codon-aware CDS alignments using PAL2NAL v.14 (Suyama et al. 2006), providing combined information from MUSCLE protein alignments (Edgar 2004) and corresponding nucleotide sequences. We retained only genes having full CDS coverage in all wasp species, and calculated p-distances for each gene using the ‘ape’ package in R (Paradis et al. 2004). Specifically, we measured p-distances for *D. muliebre* (pdistDM) and *D. ferrugineum* (pdistDF) relative to a common sexual outgroup, *D. alloeum*. Although p-distance values include evolutionary changes that have occurred in *D. alloeum* and the stem lineage preceding the divergence of *D. muliebre* and *D. ferrugineum*, using *D. alloeum* as a common reference point for comparisons allows us to infer whether a greater number of mutational differences have occurred in the asexual or sexual lineage. To achieve this, we first calculated the difference between pairwise distances (pdistDM-pdistDF) for each gene, and subsequently conducted a chi-square analysis to test against a null distribution of equal numbers of genes with greater mutational differences in sexuals and asexuals ((pdistDM>pdistDF) = (pdistDM<pdistDF)).

Variation in library preparation, sequencing, and assembly methods might generate distinct artefacts in our pipeline. To reduce the likelihood of problematic variant calling in Illumina datasets, we manually inspected a subset of genes to assess whether GATK produces accurate SNP calls across dataset types (*i.e*. DNASeq *vs*. RNASeq) (Supplemental Information). The above analyses treat each gene as an independent unit, however the actual dataset might be biased towards genes contained in linkage groups with nonindependent selective pressures. We determined the assembly scaffolds represented by the genes in the dataset to assess potential biases generated by linkage (**Supplemental Information**).

### Substitution rate analyses

We compiled the codon-aware, pairwise alignments of each CDS between either *D. alloeum* and *D. muliebre* or *D. alloeum* and *D. ferrugineum* into their own respective text files. Next, we analyzed each of these files for codon substitutions using codeml in PAML v.4.8 (Yang 2007) with a two-branch tree containing *D. alloeum* and either *D. muliebre or D. ferrugineum*. We calculated one ω ratio for all branches in the tree for each gene and calculated the average nucleotide frequency at all three codon positions, assuming no clock. We then grouped each CDS by its synonymous (dS) and non-synonymous (dN) substitution rates.

### Relative rate analyses

In addition to evaluating mutational patterns for each nuclear gene individually, we performed a global analysis to characterize whether sex loss in *Diachasma* is associated with the accumulation of deleterious mutations. We combined all CDS regions with full coverage into a single concatenated dataset. Next, we used the GTR+CAT model implemented in RAxML (Stamatakis 2006) to conduct a maximum likelihood analysis in order to estimate branch lengths between sexual and asexual species. We performed three RAxML runs, using 1) all CDS sites, 2) 1^st^+2^nd^ codon positions (≈ nonsynonymous sites), and 3) 3^rd^ codon positions (≈ synonymous sites).

We performed an implementation of Tajima’s Relative Rate test in MEGA6 (Tajima 1993; Tamura et al. 2013) to test for evolutionary rate differences in *D. muliebre* and *D. ferrugineum*. The test requires a three-taxon sequence dataset with knowledge *a priori* of the outgroup relative to the other two. Life history, morphology, and genetic data all support *D. ferrugineum* as the closest sexual relative to *D. muliebre*, with *D. alloeum* as the outgroup (Wharton and Marsh 1978; Forbes et al. 2009; Hamerlinck et al. 2016). We calculated the number of observed unique SNPs in the concatenated dataset for each *Diachasma* species. Next, a chi-square test compares the observed number of unique differences to the expected number, given equal evolutionary rates. We performed three separate analyses to test rate differences across all sites as well as approximate nonsynonymous (1^st^+2^nd^ codon positions) and synonymous (3^rd^ codon positions) sites. To be conservative, we applied the *D. muliebre* false-positives rate to the concatenated dataset to compare wasp genomes (the application of the false negatives rate as a proportion of true negatives in this case is negligible given the total length of the alignments). If ~108 (1.89%) *D. muliebre* SNPs are false positives, this individual would have 5,634 SNPs in the concatenated dataset. Applying this conservative measure, the number of unique differences was significantly greater than the 4,148 observed in *D. ferrugineum* (χ^2^ = 1416, p < 0.005).

### Intraspecific comparisons

To assess whether there is substantial variation in the nuclear genome mutation load among individual asexual females, we sequenced DNASeq datasets from five *D. muliebre* females. Applying the methods described above, we retrieved a subset of the conserved genes that had full-length coverage in the single *D. ferrugineum* female and all *D. muliebre* females. After calculating pairwise distances for each wasp relative to *D. alloeum*, we performed multiple t-tests to test for differences among the asexual distributions, applying a Bonferroni correction for multiple comparisons.

### Codon usage bias and gBGC

We conducted two separate tests to evaluate differences in genome-wide codon usage between *D. ferrugineum* and *D. muliebre* females. First, we determined the effective number of codons (ENC) for each gene (Wright 1990). The range of ENC values reflect codon usage for each amino acid, from 20 (each amino acid encoded by one codon) to 61 (all codons used equally). We calculated ENC using the program ENCprime (https://github.com/jnovembre/ENCprime), which accounts for background nucleotide compositions to enable cross-species comparisons (Novembre 2002). Second, we used the Composition Analysis Toolkit (CAT; http://www.cbrc.kaust.edu.sa/CAT) to calculate per-gene codon deviation coefficient (CDC) values (Zhang et al. 2012). CAT uses background GC and purine content for each codon position to calculate an expected codon usage value and calculates CDC as a deviation from this expectation. CDC values range from 0 (no deviation from expectation, *i.e*. no selection on codon usage) to 1 (maximum deviation, *i.e*. effective purifying selection). To assess gBGC, we used CAT to calculate per-gene GC content at third codon positions (GC3), which are most likely to be evolving neutrally. Higher GC3 values reflect preferential conversion of A and T alleles to G and C.

### Statistical analyses

For comparisons of pairwise distances, ENC, CDC, and GC3 in the genome, we performed statistical tests to determine whether the distribution of values involving *D. muliebre* and *D. ferrugineum* were different. We tested each dataset for normality using the Shapiro-Wilk method. Since all tests failed tests of normality (**Table S8**), we conducted a Wilcoxon rank sum test as a conservative measure to determine whether or not sexual and asexual wasps possess similar pairwise distance distributions.

## Supplemental Information

Supplementary Material. Figures S1 – S5. Tables S1 – S8.

