## Supplementary material for "Genome evolution in a putatively asexual wasp": Supplemtal Information

for

### Supplementary Material

#### Natural history of *Diachasma*

Wasps in genus *Diachasma* are idiobiont parasitoids of fruit flies in genus *Rhagoletis* (Wharton and Marsh 1978). Three species are known from North America: the sexual *Diachasma alloeum*, an eastern species that attacks flies in the *Rhagoletis pomonella* species complex; a second sexual species, *Diachasma ferrugineum*, which attacks the Eastern cherry fruit fly (*Rhagoletis cingulata*) in black cherry (*Prunus serotina*); and the putative asexual *Diachasma muliebre*, which attacks Western cherry fruit fly (*Rhagoeltis indifferens*) in bitter cherry (*Prunus emarginata*) and introduced sweet cherry (*Prunus avium*) (Wharton and Marsh 1978; Forbes et al. 2010).

Adult *Diachasma* wasps orient towards trees using fruit volatile cues and alight on fruits (Stelinski and Liburd 2005; Forbes et al. 2009). Females antennate the fruit surface to locate *Rhagoletis* larvae feeding inside. Upon sensing a larva, the female wasp inserts her ovipositor into the fruit and pierces the larva, laying one or more eggs. The developing wasp larva (no more than one survives per host) consumes the fly only after the fly has exited the fruit and pupated in the soil beneath the tree. Wasps overwinter in their host puparia and emerge the following year when fruits are once again ripe, completing their life cycle (Forbes et al. 2009).

We describe *D. muliebre* in this study as a *putatively* asexual species. A long history of rearing records have uncovered only female wasps across a wide geographic distribution of sites (**Table S1**). However, shortly before the completion of this study, two male wasps – apparently the first ever for the species – were discovered by A.A.F. among a collection of 510 *D. muliebre* individuals reared from a single site by Dr. Wee Yee of the USDA-ARS. To ascertain whether these males could be representatives of an undiscovered introduction of one of the sexual

*Diachasma* species found in the eastern U.S., we extracted DNA from the right legs of one of these males and then amplified and sequenced a 648 bp segment of mtCOI. This sequence proved identical to one of the five *D. muliebre* mtCOI haplotypes reported previously (6). Both males also had morphological features that matched the description of *D. muliebre* (1). Unknowns regarding these rare male *D. muliebre* are myriad, and include: Are they functional males? Does mating occur in nature and result in viable progeny? Are they haploid or diploid? How often and under what conditions are males produced? The current state of knowledge is thus that *D. muliebre* is either an asexual species that engages in rare sex, or that it is fully asexual but occasionally produces rare, but non-functional, males.

**Phylogeny of *Diachasma* and the origin of *D. muliebre* asexuality**

Morphology, host associations, and genetics all demonstrate that *D. ferrugineum* is the closest sexual relative to *D. muliebre*. These two wasps differ morphologically only in their size and the texturing on their ventral abdomen, while the other North American species, *D. alloeum*, has an obviously longer abdomen, more sculpturing differences, and several more antennal segments than either of the other two wasps (Wharton and Marsh 1978). *Diachasma muliebre* and *D. ferrugineum* also attack sister species of fly whose native host plants are both in the flowering cherry subsection of cherries (*Pseudocerasus*) (Bortiri et al. 2001), and are found on the Pacific slope and Eastern North America, respectively, suggesting different histories of post-glacial colonization. Finally, previous phylogenetic work using mitochondrial and nuclear sequence suggests that *D. muliebre* and *D. ferrugineum* are sister (Forbes et al. 2013; Hamerlinck et al. 2016), and these inferences are echoed by this study.

Previous genetic work suggests a single origin of asexuality for *D. muliebre*. Five mitochondrial haplotypes have thus far been identified, all apparently derived from a single ancestral haplotype

(Forbes et al. 2013). Further, allelic richness at five microsatellite loci genotyped for 140 individuals across all five haplotype lineages never exceeds two alleles per locus, strongly suggesting a single common ancestor that was heterozygous at many loci, and with derived lineages subsequently either maintaining or losing that heterozygosity.

All estimates of the age of the most recent common ancestor for *D. muliebre* and *D. ferrugineum* place it at less than 1 MYA. A previous study applied insect molecular clocks to *Diachasma* partial mtCOI sequences to date the split between *D. muliebre* and *D. ferrugineum* to 0.62-0.96 MYA (Forbes et al. 2013). MtCOI sequences downloaded from NCBI show differences of just 1-4 bp between *R. indifferens* and *R. cingulata* host fly species (0.11-0.44 MYA, based on molecular clocks). The lack of overlap between fly and wasp estimates suggests any of a few options: 1) that flies diverged more recently than their wasp parasites, 2) incomplete lineage sorting in the fly hosts, or 3) molecular clocks are problematic. In any case, as wasps (and their fly hosts) experience just a single generation per year, applying the most conservatively wide range of dates implies that just 110,000-960,000 generations have transpired since their split (and the inferred origin of asexuality *D. muliebre*).

**Phenotypic Evolution and Niche Expansion in *D. muliebre***

Timing of adult emergence is a trait critical to *Diachasma* wasp fitness. Adult wasps live for ~13 days (Forbes et al. 2009), during which time they must find and oviposit into *Rhagoletis* larvae within ripening fruit. Wasps that emerge too long before or any time after the temporal window of host availability will produce no progeny. In sexual *Diachasma*, adult emergence timing varies around a mean, with most wasps emerging during the height of host availability (Forbes et al. 2009; Hood et al. 2015). For *D. muliebre*, individual lineages differ in their mean emergence timing, and have more limited variation than their sexual counterparts, suggesting that the evolution of phenotypic variation among lineages allows for specialization of lineages on different temporal windows (Forbes et al. 2013). This temporal specialization of lineages may predispose *D. muliebre* for expansion into other niches. Sweet cherries (*Prunus avium*), introduced to the northwestern U.S. in 1847 (McClintock 1967) have been colonized by *R. indifferens* flies. Because sweet cherry trees set their fruit an average of 3-4 weeks earlier than bitter cherry trees, the *R. indifferens* flies in sweet cherries represent a new temporal "island" for *D. muliebre*. Only the earliest-emerging lineages of *D. muliebre* found in bitter cherries have expanded into the new sweet cherry environment (Forbes et al. 2013).

**Robust variant calling performance across Illumina sequencing datasets**

We focused on CDS comparisons in this study given that these regions are sequenced in both DNASeq and RNASeq datasets. To reduce the likelihood of problematic variant calling in NGS datasets, we performed multiple QC measures, including a) removing multi-copy genes from the dataset, b) removing genes with incomplete CDS coverage, and c) manually inspected a subset of genes to assess whether GATK produces accurate SNP calls across Illumina dataset types (DNASeq and RNASeq) and read mappers (Bowtie2 and TopHat2). We focused on genes that showed the largest absolute differences in pairwise distance values, | pdistDM – pdistDF |, as these genes could represent regions showing the greatest number of mutational differences in our dataset or alternatively could be the consequence of spurious mapping.

By analyzing 50 genes with the highest pdistDM value relative to pdistDF **(Table S5)**, we identified 477 SNPs in *D. muliebre* reads mapped with Bowtie2, nine (1.89%) of which were determined to be false positives and were contained on a single gene. We confirmed 146 SNPs in *D. ferrugineum* in these same genes mapped with TopHat2; zero were false positives. Additionally, we identified zero and three SNPs that were false negatives in *D. muliebre* and *D.*

*ferrugineum*, respectively. In the 50 genes with the highest pdistDF values relative to pdistDM (**Table S6**), we identified 259 SNPs in *D. ferrugineum* and 111 SNPs in *D. muliebre*, none of which were false positives. Two genes contained false negative SNPs in *D. muliebre*, however one was a part of a tandem duplication and was subsequently removed from the dataset. Overall, our SNP calling pipeline displayed consistent performance across dataset types (DNASeq and RNASeq) and read mappers (Bowtie2 and TopHat2).

**Low levels of intraspecific variation in *Diachasma***

Illumina DNASeq data from five *D. muliebre* females mapped to 1,764 complete CDS regions. Consistent with comparisons between *D. ferrugineum* and a single *D. muliebre* haplotype, mean pairwise distances in all *D. muliebre* haplotypes were larger than the mean observed in *D. ferrugineum* (**Table S4**). This pattern can be visualized as a rightward shift in the mutation landscape histograms (**Figure S1**). In contrast, pairwise distances between pairs of *D. muliebre* females were not statistically significant (10 comparisons, Wilcoxon rank sum $p > 0.005$ with Bonferroni correction), providing no evidence for variation in mutation accumulation landscapes within asexuals (**Table S4**, **Figure S2**). Using maximum likelihood analyses of all codon positions, *D. muliebre* haplotype branches were small relative to the stem branch leading up to the common ancestor of all *D. muliebre* individuals (**Figure S3**). The observed excess of nucleotide changes in the *D. muliebre* lineage is consistent with the pattern observed in the larger gene dataset that included a single *D. muliebre* individual.

The total length of CDS regions with >2X coverage in sexual and asexual *Diachasma* females was 734,668 bp. At these sites, *D. muliebre* females had a consistently higher number of total SNPs relative to *D. ferrugineum* (**Table S7**). The percentage of variant sites in these regions (0.00546 in *D. muliebre* I, 0.00496 in *D. ferrugineum* RNA) were similar to the mean pairwise distance values obtained in the full CDS dataset (0.005496 in *D. muliebre* I, 0.005089 in *D. ferrugineum* RNA). Of the total SNPs, ~1% were heterozygous in *D. muliebre* individuals, a considerably smaller percentage than found in *D. ferrugineum* (~18-20%), supporting that asexual reproduction in *Diachasma* is accompanied by a reduction in genomic heterozygosity (**Table S7**).

**No apparent linkage biases in pairwise distance measures**

To assess biases in the genome locations of sampled genes, we determined the assembly scaffold of each gene in the dataset. Next, we counted (a) the scaffolds sampled by 10% of genes with largest absolute differences in pairwise distance values, and (b) the subdivisions of scaffolds in (a) by the largest pairwise distance values in either *D. ferrugineum* or *D. muliebre*. If there are strong biases in the dataset, the scaffolds contained in the subdivided datasets in (b) should be distinct pools of scaffolds from the assembly.

To address this, we quantified the number of scaffolds sampled in the entire gene dataset, as well as the scaffolds containing genes with divergent molecular evolution in *D. ferrugineum* and *D. muliebre*. Of 3,968 total scaffolds in the *D. alloeum* assembly, 274 (7.42%) contained the 3,127 genes used for pairwise comparisons. The low percentage is unsurprising given that we restricted our gene set to those conserved between *D. alloeum* and two other braconid wasps, which are most likely contained on a small number of long scaffolds in the assembly. A subset of genes with the highest pairwise distances in *D. muliebre* relative to *D. ferrugineum* (313 genes, ~10%) were contained in 102 scaffolds, whereas a subset of genes with the reverse pattern (highest in *D. ferrugineum*) were contained in 121 scaffolds. Finally, the combined set of 626 genes were on 141 scaffolds, providing evidence that the subsets of scaffolds at the ends of the distribution are largely overlapping. In other words, many scaffolds likely contain genes showing either accelerated evolution in *D. ferrugineum* or *D. muliebre*, and the acceleration in molecular evolution in the asexual wasp species is not explained by selection acting at linked sites.

**No evidence of codon usage bias in *Diachasma***

ENC and CDC values were comparable in *D. ferrugineum* and *D. muliebre* (Wilcoxon rank sum $p > 0.9$ in all cases) when analyzing the 3,127 genes with full sequences in two wasp species, indicating similar codon usage biases across reproductive modes (**Table S3**). The results suggest that accelerated evolution in *D. muliebre* is not explained by relaxed selection on codon usage. In addition, per-gene GC3 distributions were not significantly different in *D. ferrugineum* and *D. muliebre* (Wilcoxon rank sum $p \geq 0.47$), suggesting that gBGC activity is unchanged in asexual wasps and is not contributing to their increased mutational load.

### Supplementary Figures

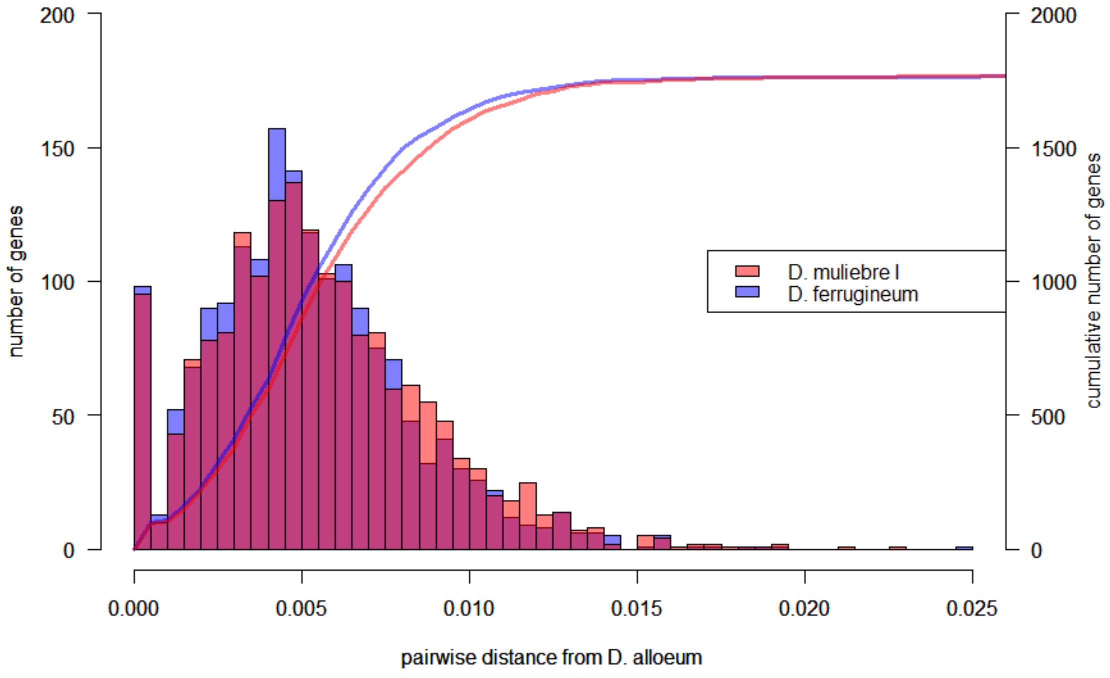

**Figure S1. Mutation landscapes of sexual and various asexual *Diachasma* wasps.**

Columns represent binned frequencies (left y-axis) of corresponding p-distance values from *D. alloeum*. Curves represent the cumulative frequency of binned columns (right y-axis).

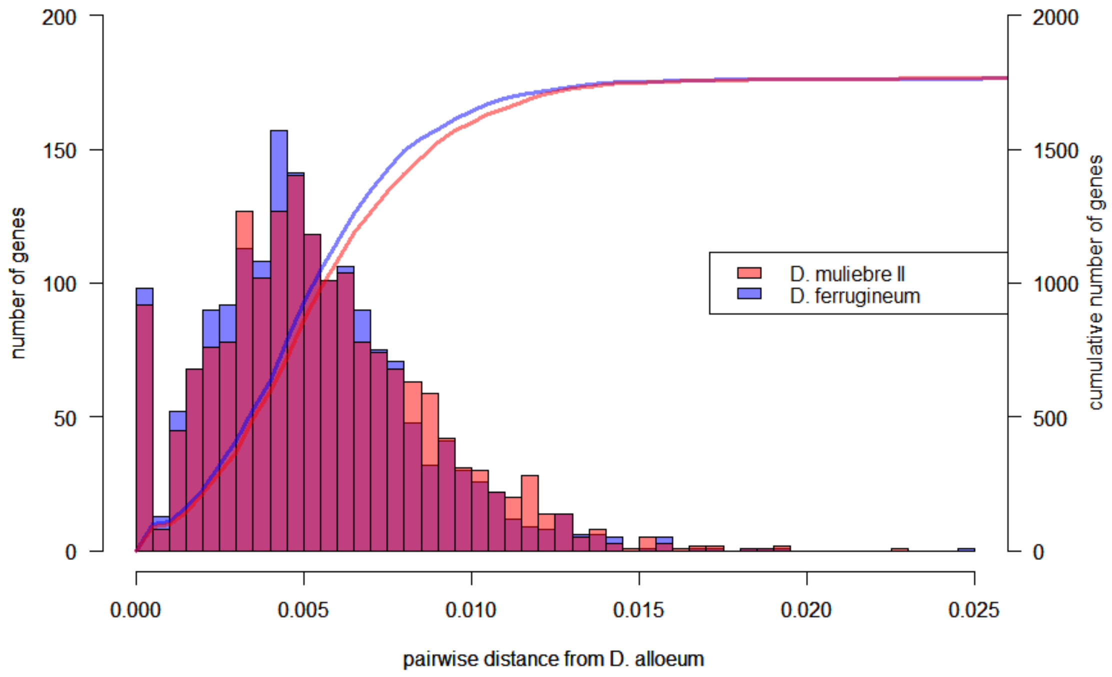

**Figure S1 – continued**

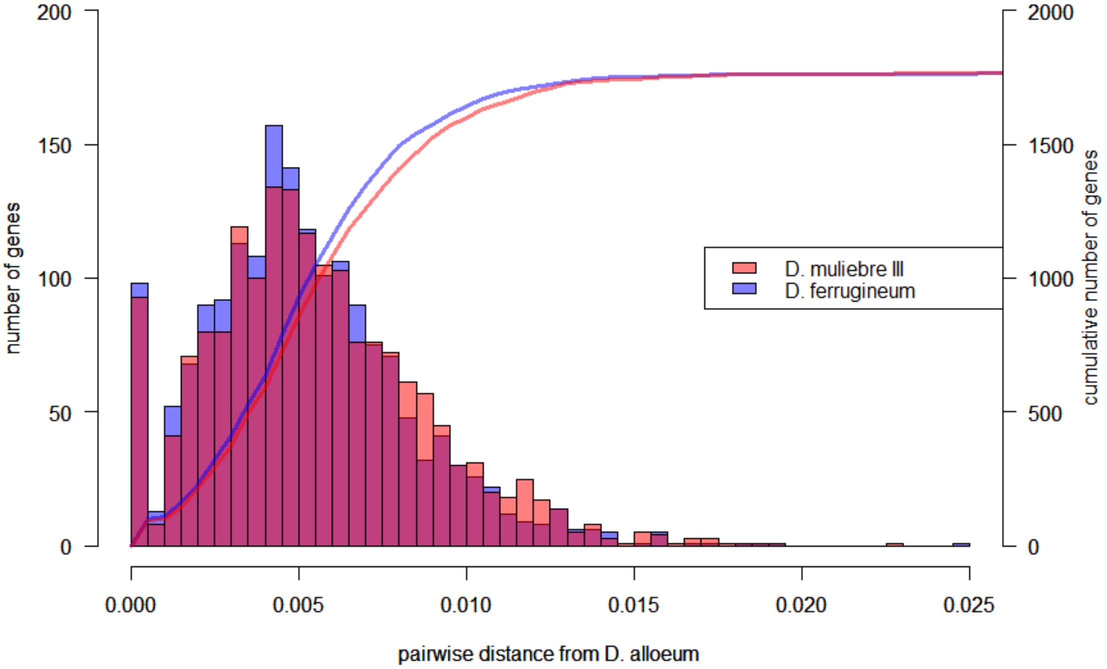

**Figure S1 – continued**

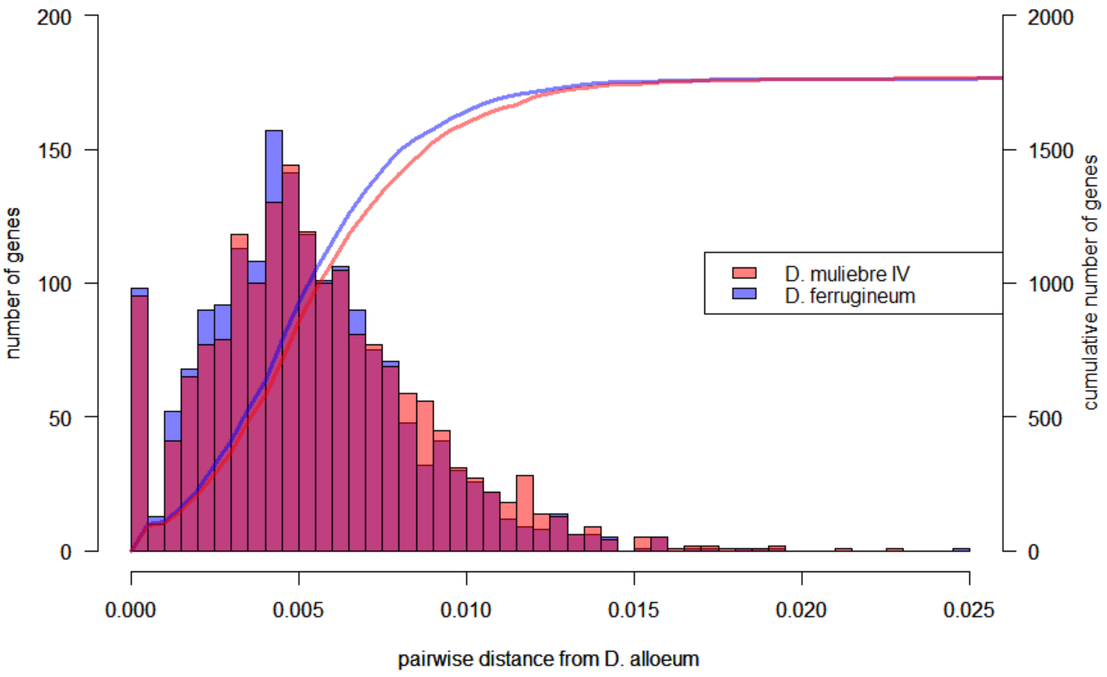

**Figure S1 – continued**

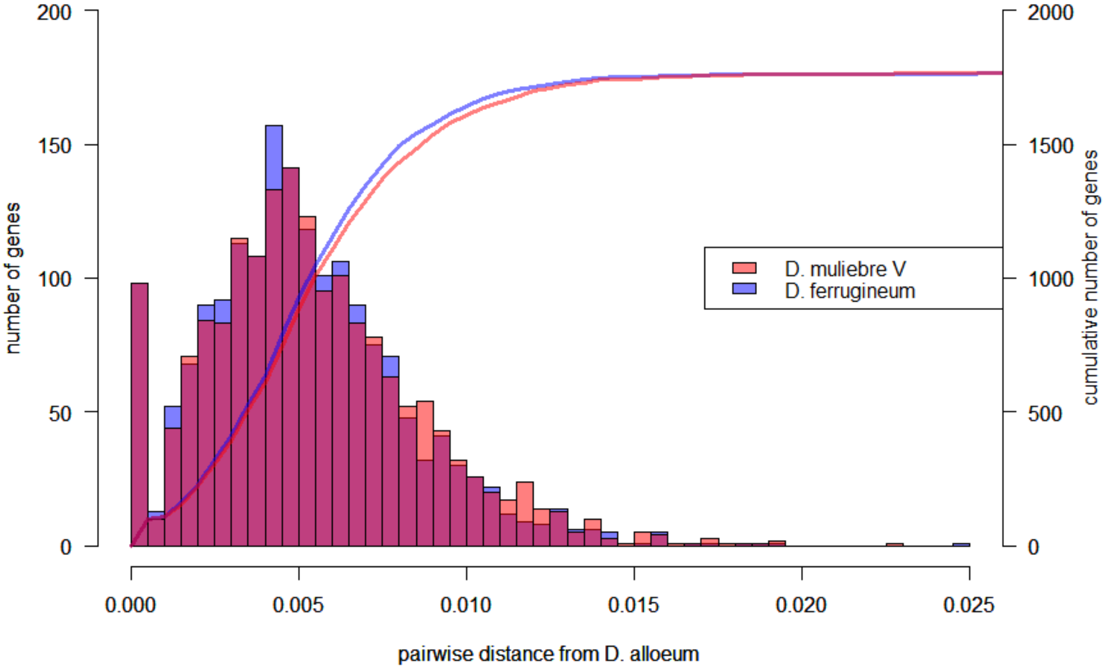

**Figure S1 – continued**

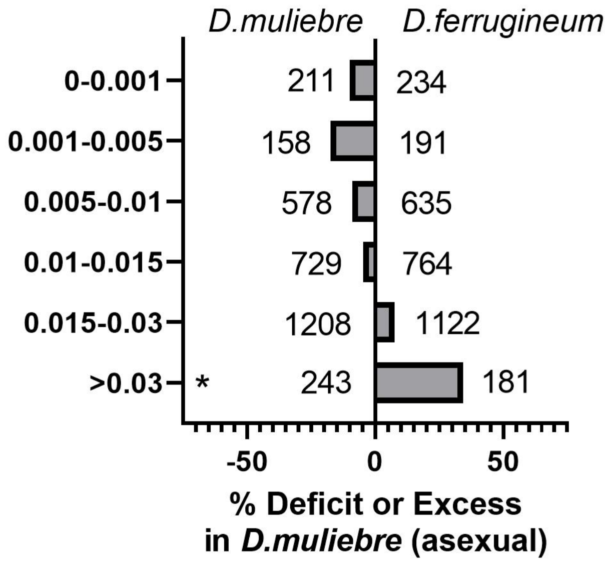

**Figure S2. Comparison of silent substitutions (dS) in 3,127 orthologs from *D. muliebre* and *D. ferrigineum*, compared to *D. alloeum*.**

Silent substitutions (dS) were calculated in PAML (ref) comparing *D. muliebre* and *D. ferrigineum* to *D. alloeum* in three species alignments. The total number of genes in each bin for each species is shown.

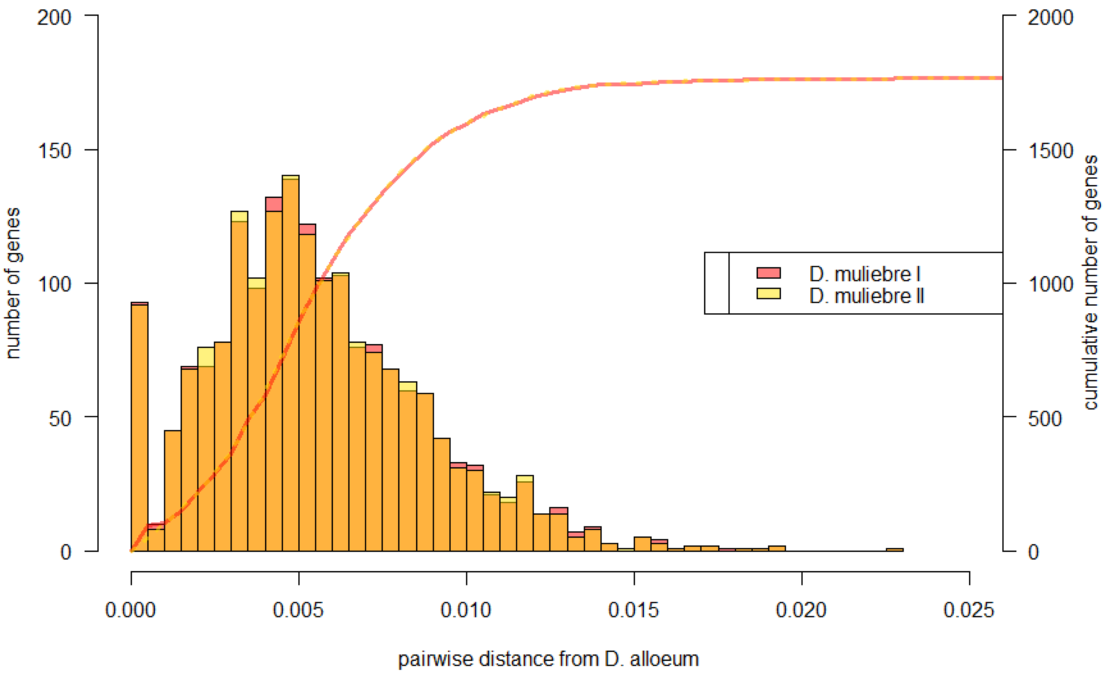

**Figure S3. Mutation landscapes of various asexual *Diachasma* wasps.**

Columns represent binned frequencies (left y-axis) of corresponding p-distance values from *D. alloeum*. Curves represent the cumulative frequency of binned columns (right y-axis).

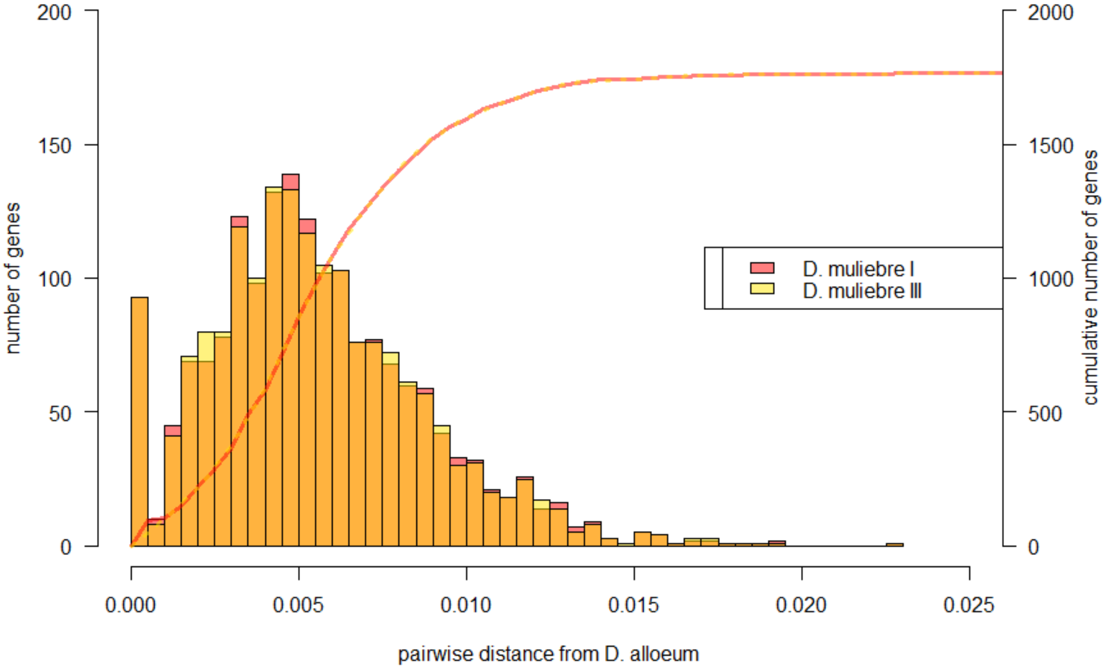

**Figure S3 – continued**

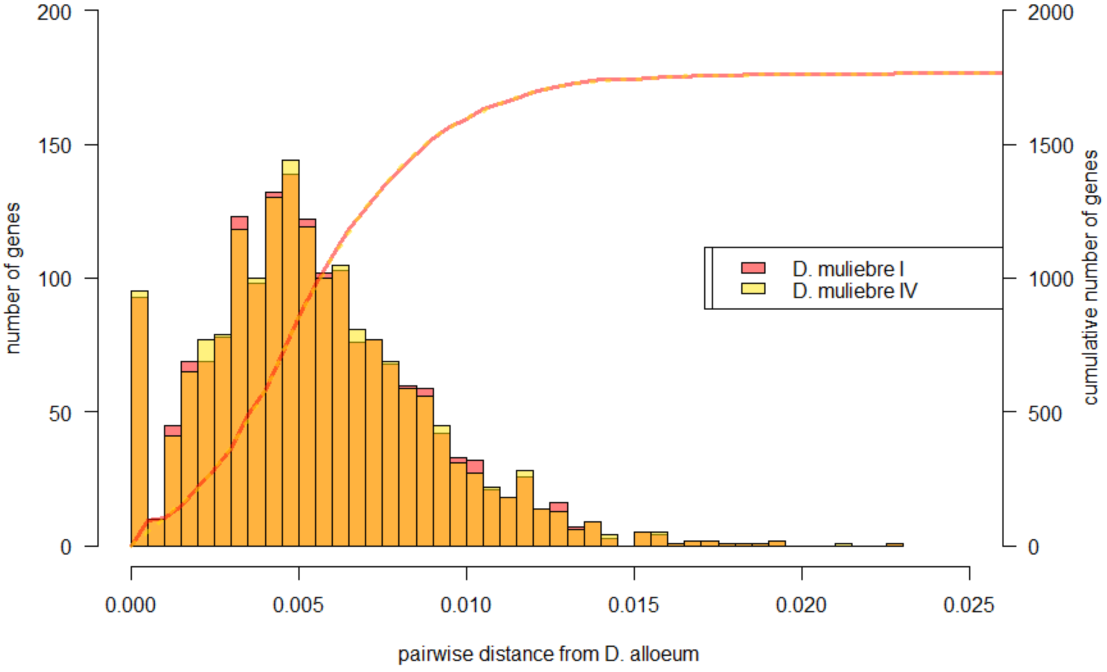

**Figure S3 – continued**

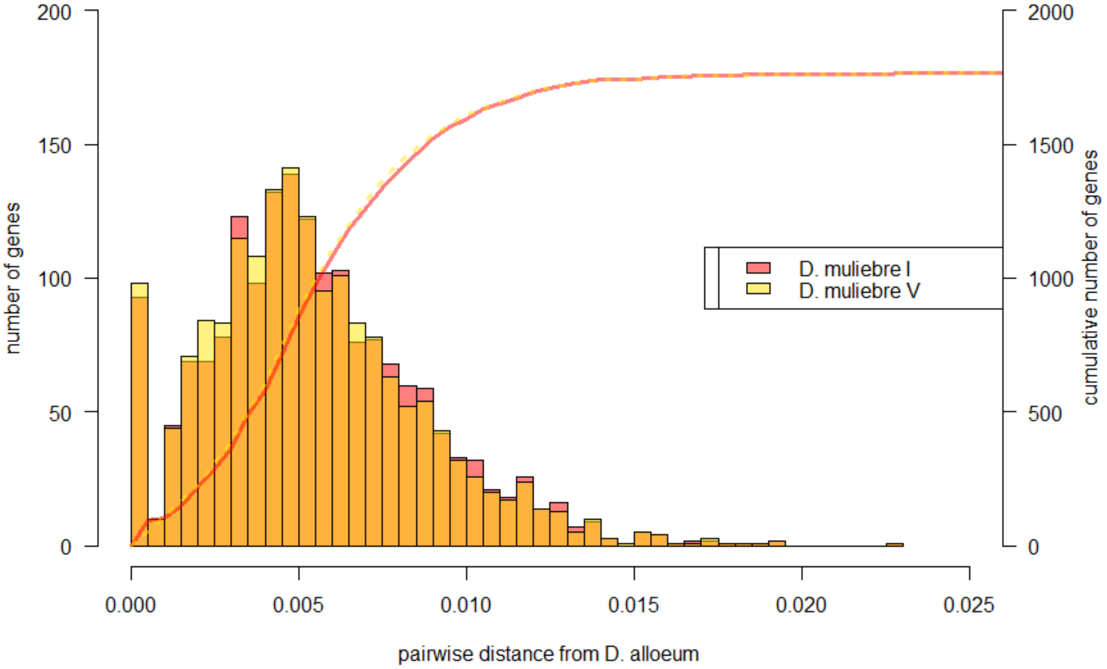

**Figure S3 – continued**

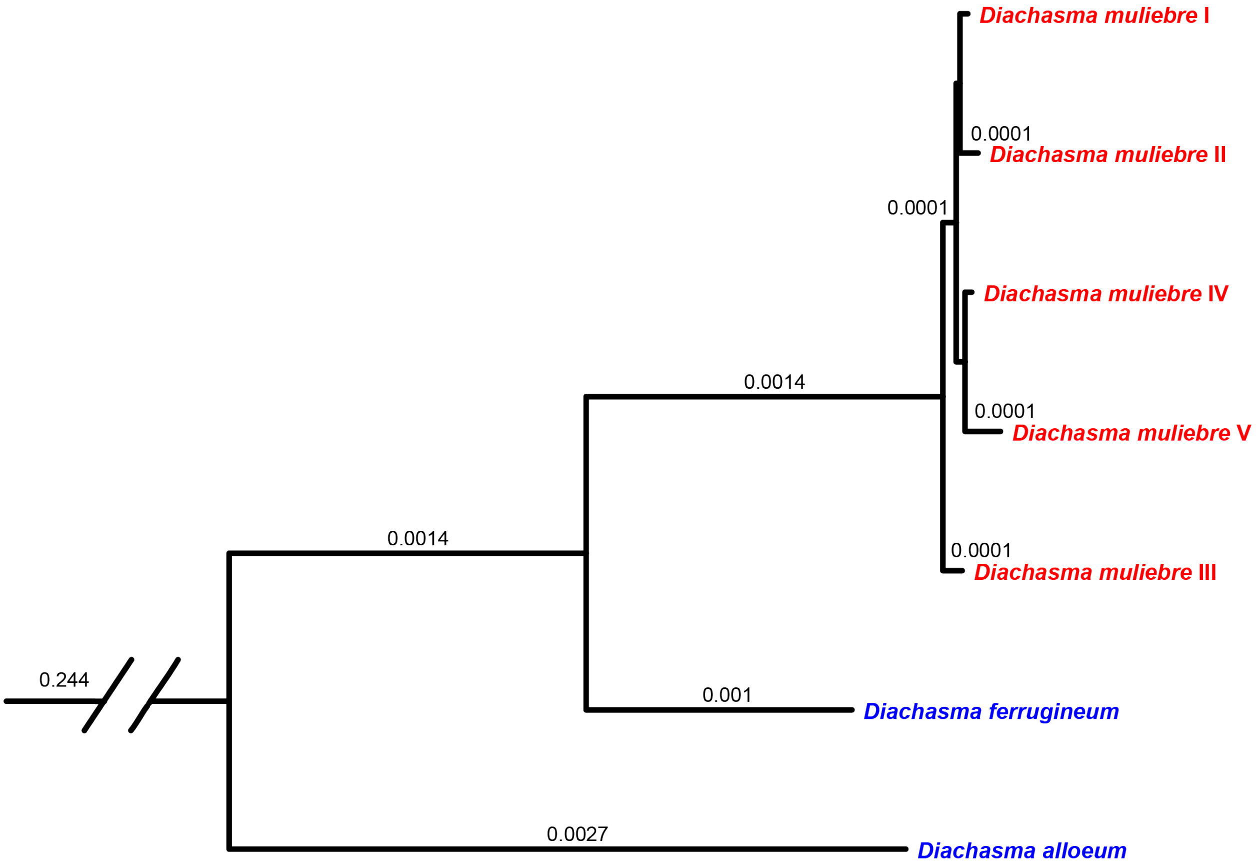

**Figure S4. Maximum likelihood analysis of concatenated nuclear gene dataset among *D. muliebre* haplotypes.**

Asexual (red) and sexual (blue) *Diachasma* lineages are shown, and branch lengths ≥ 0.0001 are labeled. Phylogenetic tree of all sites conserved in all *D. muliebre* haplotypes (1,727,505 bp). Break in tree divides outgroup (*Fopius arisanus*, not shown) to *Diachasma* clade.

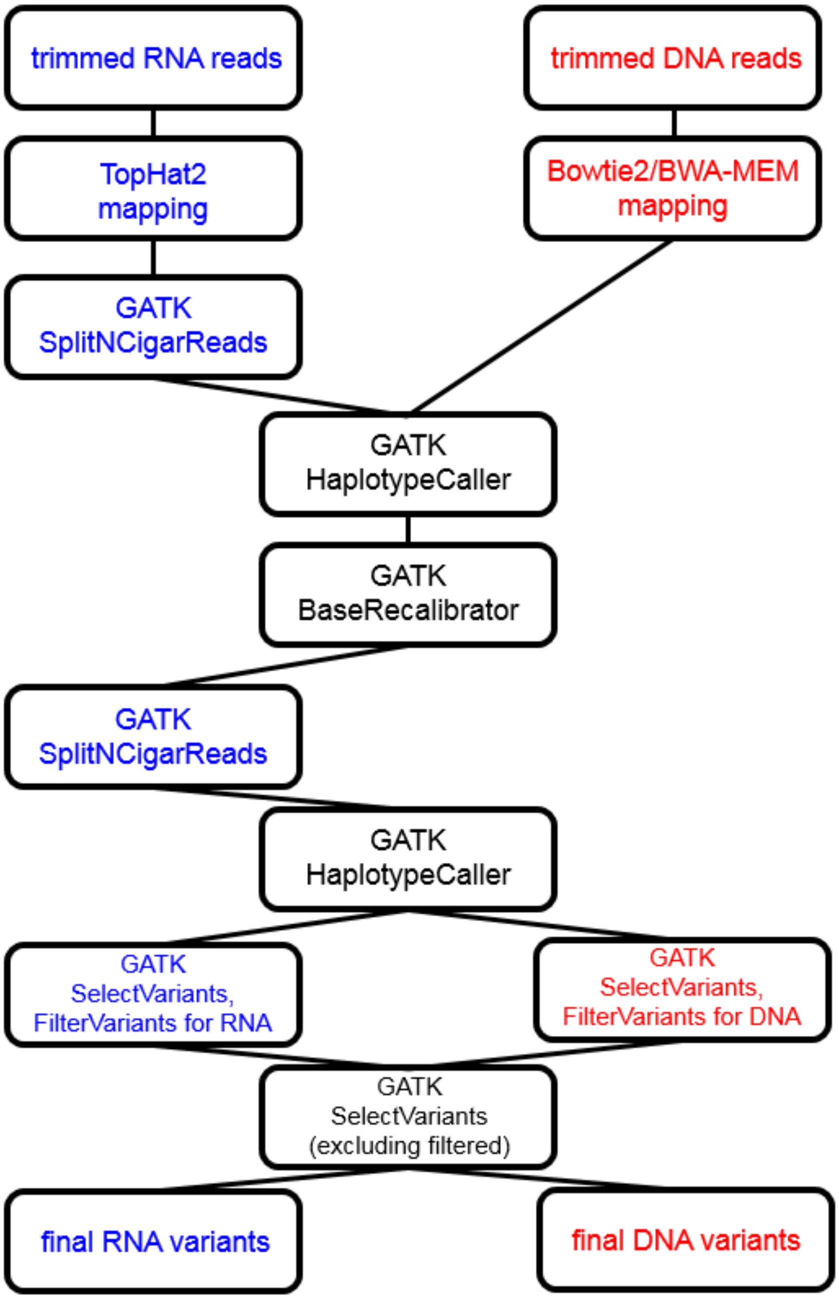

**Figure S5. Diagram of variant calling pipeline used for *Diachasma* sequencing datasets.** Blue and red text refers to steps specific for RNA (*D. ferrugineum* sexual female) and DNA (*D. muliebre* asexual female) datasets, black text refers to steps used for all datasets using identical parameters.

### Supplementary Tables

**Table S1. Sex ratios of historic *D. muliebre* collections.**

| Year | Collector | Location | Female | Male | Reference |
|---|---|---|---|---|---|
| 1954 | K.E. Frick | White Salmon, WA | 112 | 0 | Muesebeck 1956 |
| 1954 | K.E. Frick | WA, location not recorded | 147[a] | 0 | Muesebeck 1956 |
| 1954 | K.E. Frick | OR, location not recorded | | | |
| 1954 | K.E. Frick | Placer Co., CA | | | |
| 1954 | K.E. Frick | Belknap Springs, OR | | | |
| 1997 | G. Bush | Calaveras Co., CA | 11 | 0 | J. Smith, pers. comm. |
| 2010 | W. Yee | Roslyn, WA | 643 | 0 | Yee et al. 2015 |
| 2010 | W. Yee | Nile, WA | 250 | 0 | Yee et al. 2015 |
| 2011 | W. Yee | Roslyn, WA | 2 | 0 | Yee et al. 2015 |
| 2011 | W. Yee | Nile, WA | 426 | 0 | Yee et al. 2015 |
| 2011 | W. Yee | Flathead Lake, MT | 188 | 0 | Wee Yee and A.A.F., pers. obs. |
| 2012 | W. Yee | Roslyn, WA | 540 | 0 | Yee et al. 2015 |
| 2013 | W. Yee | Flathead Lake, MT | 130 | 0 | Wee Yee and A.A.F., pers. obs. |
| 2020 | W. Yee | Nile, WA | 508 | 2[b] | W. Yee, unpublished |
| 2020 | W. Yee | Roslyn, WA | | | |
| 2020 | W. Yee | Cle Elum, WA | | | |

[a] Muesebeck (1956) did not specify how many of these 147 *D. muliebre* were collected at each of four locations.

[b] The 510 wasps from 2020 were collected from three locations in Washington but the exact distribution of wasps across the three sites has not yet been tallied and will be reported in a future paper. Most wasps (>60%), including both of the male *D. muliebre*, were from the Roslyn, WA site.

**Table S2. Summary of Illumina sequencing datasets.**

| wasp sample | Library type | Library design | Read count (pairs) | SRA accession |
|---|---|---|---|---|
| D. muliebre I | DNASeq | 2 x 300 | 15,417,069 | TBD |
| D. muliebre II | DNASeq | 2 x 300 | 14,041,561 | TBD |
| D. muliebre III | DNASeq | 2 x 300 | 15,229,838 | TBD |
| D. muliebre IV | DNASeq | 2 x 300 | 12,645,301 | TBD |
| D. muliebre V | DNASeq | 2 x 300 | 16,226,907 | TBD |
| D. ferrugineum RNA | RNASeq | 2 x 300 | 8,313,856 | TBD |
| D. ferrugineum RRL | DNASeq | 2 x 250 | 2,211,235 | TBD |

**Table S3. Measures of codon bias and nucleotide composition in *Diachasma*.**

| | D. ferrugineum mean (SD) | D. muliebre I mean (SD) | # genes | W-statistic | p-value |
|---|---|---|---|---|---|
| ENC | 54.69482 (3.69) | 54.69372 (3.70) | 3127 | 4887500 | 0.98 |
| CDC | 0.130546 (0.05) | 0.130613 (0.05) | 3127 | 4883900 | 0.94 |
| GC3 | 0.52141 (0.089) | 0.521456 (0.089) | 3127 | 4887500 | 0.98 |

ENC = effective number of codons, CDC = codon deviation coefficient, GC3 = GC content at third codon position. Wilcoxon rank sum tests were used to assess significant differences in means between *D. ferrugineum* (sexual, blue) and *D. muliebre* (asexual, red).

**Table S4. Pairwise distances in nuclear CDS regions in *Diachasma*.**

| Wasp 1<br>Mpdist (Variance) | Wasp 2<br>Mpdist (Variance) | #<br>genes | W-statistic | p-value |
|---|---|---|---|---|
| D. ferrugineum<br>0.005089 (7.95E-6) | D. muliebre I<br>0.005496 (9.36E-6) | 3127 | 4527800 | 4.17E-7 |
| D. ferrugineum<br>0.005173 (9.35E-6) | D. muliebre I<br>0.005534 (1.07E-5) | 1764 | 1461800 | 1.88E-3 |
| D. muliebre I<br>0.005534 (1.07E-5) | D. muliebre II<br>0.005508 (1.04E-5) | 1764 | 1561700 | 0.85 |
| D. muliebre I<br>0.005534 (1.07E-5) | D. muliebre III<br>0.005521 (1.06E-5) | 1764 | 1560200 | 0.89 |
| D. muliebre I<br>0.005534 (1.07E-5) | D. muliebre IV<br>0.005534 (1.08E-5) | 1764 | 1557200 | 0.96 |
| D. muliebre I<br>0.005534 (1.07E-5) | D. muliebre V<br>0.005414 (1.05E-5) | 1764 | 1591900 | 0.23 |
| D. muliebre II<br>0.005508 (1.04E-5) | D. muliebre III<br>0.005521 (1.06E-5) | 1764 | 1554300 | 0.96 |
| D. muliebre II<br>0.005508 (1.04E-5) | D. muliebre IV<br>0.005534 (1.08E-5) | 1764 | 1551300 | 0.88 |
| D. muliebre II<br>0.005508 (1.04E-5) | D. muliebre V<br>0.005414 (1.05E-5) | 1764 | 1586000 | 0.32 |
| D. muliebre III<br>0.005521 (1.06E-5) | D. muliebre IV<br>0.005534 (1.08E-5) | 1764 | 1552800 | 0.92 |

| | | | | |
|---|---|---|---|---|
| D. muliebre III<br><br>0.005521 (1.06E-5) | D. muliebre V<br><br>0.005414 (1.05E-5) | 1764 | 1587600 | 0.29 |
| D. muliebre IV<br><br>0.005534 (1.08E-5) | D. muliebre V<br><br>0.005414 (1.05E-5) | 1764 | 1590600 | 0.25 |

Wilcoxon rank sum tests were used to assess significant differences in distributions of pairwise distance measures involving *D. ferrugineum* (sexual) and *D. muliebre* (asexual) relative to a common outgroup (*D. alloeum*, sexual).

**Table S5. Manual annotation of SNPs in accelerated asexual genes.**

| coordinates | pdist DF | pdist DM | Δ pdist | DF SNP | DF FP | DF FN | DM SNP | DM FP | DM FN |
|---|---|---|---|---|---|---|---|---|---|
| NW_015145034.1: 1825605-1826552 | 0.0038 | 0.0246 | 0.0208 | 2 | 0 | 0 | 13 | 0 | 0 |
| NW_015145098.1: 55721-58954 | 0.0025 | 0.0198 | 0.0173 | 1 | 0 | 0 | 8 | 0 | 0 |
| NW_015145106.1: 881760-883391 | 0.0023 | 0.0159 | 0.0136 | 1 | 0 | 0 | 7 | 0 | 0 |
| NW_015145106.1: 878338-879847 | 0.0000 | 0.0132 | 0.0132 | 0 | 0 | 0 | 5 | 0 | 0 |
| NW_015145163.1: 888976-900771 | 0.0007 | 0.0136 | 0.0129 | 1 | 0 | 0 | 19 | 0 | 0 |
| NW_015145023.1: 2281296-2282712 | 0.0064 | 0.0191 | 0.0127 | 3 | 0 | 0 | 9 | 0 | 0 |

**Table S5 – continued**

| | | | | | | | | | |
|---|---|---|---|---|---|---|---|---|---|
| NW_015145157.1: 143688-146013 | 0.0062 | 0.0186 | 0.0124 | 3 | 0 | 0 | 9 | 0 | 0 |
| NW_015145005.1: 5733031-5735567 | 0.0031 | 0.0154 | 0.0123 | 3 | 0 | 0 | 15 | 0 | 0 |
| NW_015145037.1: 578137-579988 | 0.0000 | 0.0119 | 0.0119 | 0 | 0 | 0 | 2 | 0 | 0 |
| NW_015145153.1: 945274-947100 | 0.0056 | 0.0167 | 0.0111 | 2 | 0 | 0 | 6 | 0 | 0 |
| NW_015145095.1: 97674-103675 | 0.0044 | 0.0153 | 0.0109 | 2 | 0 | 0 | 7 | 0 | 0 |
| NW_015145025.1: 2434869-2442720 | 0.0035 | 0.0139 | 0.0104 | 4 | 0 | 0 | 16 | 0 | 0 |
| NW_015145003.1: 656878-660440 | 0.0000 | 0.0100 | 0.0100 | 0 | 0 | 0 | 4 | 0 | 0 |

**Table S5 – continued**

| | | | | | | | | | |
|---|---|---|---|---|---|---|---|---|---|
| NW_015145023.1: 2133414-2143862 | 0.0010 | 0.0104 | 0.0094 | 1 | 0 | 0 | 10 | 0 | 0 |
| NW_015148955.1: 1184829-1186954 | 0.0037 | 0.0130 | 0.0093 | 2 | 0 | 0 | 7 | 0 | 0 |
| NW_015145003.1: 804104-806929 | 0.0013 | 0.0104 | 0.0091 | 1 | 0 | 1 | 8 | 0 | 0 |
| NW_015145005.1: 5504884-5508136 | 0.0000 | 0.0091 | 0.0091 | 0 | 0 | 0 | 4 | 0 | 0 |
| NW_015145178.1: 338673-340502 | 0.0057 | 0.0144 | 0.0086 | 2 | 0 | 0 | 5 | 0 | 0 |
| NW_015145178.1: 258040-261337 | 0.0028 | 0.0114 | 0.0085 | 2 | 0 | 0 | 8 | 0 | 0 |
| NW_015145020.1: 45267-46162 | 0.0028 | 0.0112 | 0.0084 | 1 | 0 | 0 | 4 | 0 | 0 |

**Table S5 – continued**

| | | | | | | | | | |
|---|---|---|---|---|---|---|---|---|---|
| NW_015145005.1: 4120625-4125899 | 0.0035 | 0.0118 | 0.0084 | 5 | 0 | 1 | 17 | 9 | 0 |
| NW_015145350.1: 251188-253471 | 0.0083 | 0.0167 | 0.0083 | 2 | 0 | 1 | 4 | 0 | 0 |
| NW_015145163.1: 227785-230673 | 0.0042 | 0.0125 | 0.0083 | 3 | 0 | 0 | 9 | 0 | 0 |
| NW_015145350.1: 59677-62423 | 0.0081 | 0.0163 | 0.0081 | 4 | 0 | 0 | 8 | 0 | 0 |
| NW_015145320.1: 72756-187265 | 0.0016 | 0.0097 | 0.0081 | 1 | 0 | 0 | 6 | 0 | 0 |
| NW_015145163.1: 915030-919136 | 0.0010 | 0.0088 | 0.0078 | 1 | 0 | 0 | 9 | 0 | 0 |
| NW_015145099.1: 944458-951709 | 0.0048 | 0.0126 | 0.0078 | 5 | 0 | 0 | 13 | 0 | 0 |

**Table S5 – continued**

| | | | | | | | | | |
|---|---|---|---|---|---|---|---|---|---|
| NW_015145615.1: 161037-165080 | 0.0034 | 0.0112 | 0.0077 | 4 | 0 | 0 | 13 | 0 | 0 |
| NW_015145006.1: 875190-883971 | 0.0000 | 0.0074 | 0.0074 | 0 | 0 | 0 | 2 | 0 | 0 |
| NW_015148955.1: 1513657-1517544 | 0.0041 | 0.0115 | 0.0074 | 5 | 0 | 0 | 14 | 0 | 0 |
| NW_015148955.1: 81090-84019 | 0.0027 | 0.0099 | 0.0072 | 3 | 0 | 0 | 11 | 0 | 0 |
| NW_015145006.1: 1150941-1156578 | 0.0051 | 0.0123 | 0.0072 | 5 | 0 | 0 | 12 | 0 | 0 |
| NW_015145005.1: 2102771-2104667 | 0.0054 | 0.0125 | 0.0072 | 3 | 0 | 0 | 7 | 0 | 0 |
| NW_015145005.1: 4595569-4597668 | 0.0048 | 0.0119 | 0.0071 | 2 | 0 | 0 | 5 | 0 | 0 |

**Table S5 – continued**

| | | | | | | | | | |
|---|---|---|---|---|---|---|---|---|---|
| NW_015145350.1: 208163-209673 | 0.0080 | 0.0152 | 0.0071 | 9 | 0 | 0 | 17 | 0 | 0 |
| NW_015145161.1: 420508-422178 | 0.0047 | 0.0118 | 0.0071 | 2 | 0 | 0 | 5 | 0 | 0 |
| NW_015145070.1: 2316872-2318049 | 0.0000 | 0.0070 | 0.0070 | 0 | 0 | 0 | 4 | 0 | 0 |
| NW_015145152.1: 6995-28417 | 0.0000 | 0.0069 | 0.0069 | 0 | 0 | 0 | 5 | 0 | 0 |
| NW_015145023.1: 1617565-1626985 | 0.0011 | 0.0080 | 0.0068 | 1 | 0 | 0 | 7 | 0 | 0 |
| NW_015145157.1: 136500-138970 | 0.0049 | 0.0117 | 0.0068 | 5 | 0 | 0 | 12 | 0 | 0 |
| NW_015145860.1: 255652-257517 | 0.0060 | 0.0128 | 0.0068 | 8 | 0 | 0 | 17 | 0 | 0 |

**Table S5 – continued**

| | | | | | | | | | |
|---|---|---|---|---|---|---|---|---|---|
| NW_015145098.1: 988521-996880 | 0.0061 | 0.0129 | 0.0068 | 9 | 0 | 0 | 19 | 0 | 0 |
| NW_015145661.1: 400051-401741 | 0.0034 | 0.0102 | 0.0068 | 2 | 0 | 0 | 6 | 0 | 0 |
| NW_015145533.1: 149632-211538 | 0.0048 | 0.0116 | 0.0068 | 10 | 0 | 0 | 24 | 0 | 0 |
| NW_015145075.1: 1883704-1885561 | 0.0101 | 0.0168 | 0.0067 | 9 | 0 | 0 | 15 | 0 | 0 |
| NW_015145004.1: 2078622-2083845 | 0.0044 | 0.0111 | 0.0066 | 4 | 0 | 0 | 10 | 0 | 0 |
| NW_015145229.1: 965720-967127 | 0.0013 | 0.0079 | 0.0066 | 1 | 0 | 0 | 6 | 0 | 0 |
| NW_015145098.1: 63162-69263 | 0.0022 | 0.0087 | 0.0065 | 1 | 0 | 0 | 4 | 0 | 0 |

**Table S5 – continued**

| | | | | | | | | | |
|---|---|---|---|---|---|---|---|---|---|
| NW_015145009.1: 1652636-1658501 | 0.0050 | 0.0115 | 0.0065 | 7 | 0 | 0 | 16 | 0 | 0 |
| NW_015145079.1: 384157-404721 | 0.0026 | 0.0090 | 0.0064 | 4 | 0 | 0 | 14 | 0 | 0 |
| **Total** | | | | 146 | 0 | 3 | 477 | 9 | 0 |

FP = False Positive, FN = False Negative.

**Table S6. Manual annotation of SNPs in accelerated sexual genes.**

| coordinates | pdist DF | pdist DM | Δ pdist | DF SNP | DF FP | DF FN | DM SNP | DM FP | DM FN |
|---|---|---|---|---|---|---|---|---|---|
| NW_015145088.1: 401730-402966 | 0.0126 | 0.0000 | -0.0126 | 6 | 0 | 0 | 0 | 0 | 0 |
| NW_015145025.1: 909373-915643 | 0.0093 | 0.0000 | -0.0093 | 2 | 0 | 0 | 0 | 0 | 0 |
| NW_015145068.1: 991747-995522 | 0.0150 | 0.0075 | -0.0075 | 8 | 0 | 0 | 4 | 0 | 0 |
| NW_015145004.1: 2083680-2085654 | 0.0075 | 0.0000 | -0.0075 | 4 | 0 | 0 | 0 | 0 | 13 |
| NW_015145036.1: 1408133-1408755 | 0.0071 | 0.0000 | -0.0071 | 2 | 0 | 0 | 0 | 0 | 0 |
| NW_015145078.1: 569758-571332 | 0.0100 | 0.0033 | -0.0067 | 3 | 0 | 0 | 1 | 0 | 0 |

**Table S6 - continued**

| | | | | | | | | | |
|---|---|---|---|---|---|---|---|---|---|
| NW_015145005.1: 4708964-4713497 | 0.0063 | 0.0000 | -0.0063 | 3 | 0 | 0 | 0 | 0 | 0 |
| NW_015145693.1: 373960-374854 | 0.0061 | 0.0000 | -0.0061 | 2 | 0 | 0 | 0 | 0 | 0 |
| NW_015145845.1: 55880-85443 | 0.0058 | 0.0000 | -0.0058 | 2 | 0 | 0 | 0 | 0 | 2 |
| NW_015145060.1: 670567-671740 | 0.0138 | 0.0083 | -0.0055 | 5 | 0 | 0 | 3 | 0 | 0 |
| NW_015145093.1: 404718-418275 | 0.0071 | 0.0016 | -0.0055 | 13 | 0 | 0 | 3 | 0 | 0 |
| NW_015145044.1: 490527-492944 | 0.0082 | 0.0027 | -0.0054 | 6 | 0 | 0 | 2 | 0 | 0 |
| NW_015145034.1: 2086488-2088029 | 0.0127 | 0.0072 | -0.0054 | 7 | 0 | 0 | 4 | 0 | 0 |

**Table S6 - continued**

| | | | | | | | | | |
|---|---|---|---|---|---|---|---|---|---|
| NW_015145137.1: 705607-709105 | 0.0065 | 0.0011 | -0.0054 | 6 | 0 | 0 | 1 | 0 | 0 |
| NW_015145095.1: 1425155-1427832 | 0.0144 | 0.0090 | -0.0054 | 8 | 0 | 0 | 5 | 0 | 0 |
| NW_015145028.1: 2104322-2114165 | 0.0053 | 0.0000 | -0.0053 | 1 | 0 | 0 | 0 | 0 | 0 |
| NW_015145332.1: 180088-182444 | 0.0092 | 0.0039 | -0.0052 | 7 | 0 | 0 | 3 | 0 | 0 |
| NW_015145025.1: 655387-658940 | 0.0065 | 0.0013 | -0.0052 | 5 | 0 | 0 | 1 | 0 | 0 |
| NW_015145006.1: 3869219-3869732 | 0.0103 | 0.0051 | -0.0051 | 2 | 0 | 0 | 1 | 0 | 0 |
| NW_015145085.1: 355742-359769 | 0.0062 | 0.0012 | -0.0050 | 5 | 0 | 0 | 1 | 0 | 0 |

**Table S6 - continued**

| | | | | | | | | | |
|---|---|---|---|---|---|---|---|---|---|
| NW_015145027.1: 202486-204481 | 0.0066 | 0.0017 | -0.0050 | 4 | 0 | 0 | 1 | 0 | 0 |
| NW_015145020.1: 772046-773302 | 0.0173 | 0.0123 | -0.0049 | 7 | 0 | 0 | 5 | 0 | 0 |
| NW_015145002.1: 2519224-2522717 | 0.0082 | 0.0033 | -0.0049 | 5 | 0 | 0 | 2 | 0 | 0 |
| NW_015145002.1: 1028020-1030852 | 0.0048 | 0.0000 | -0.0048 | 3 | 0 | 0 | 0 | 0 | 0 |
| NW_015145159.1: 244616-245469 | 0.0095 | 0.0048 | -0.0048 | 2 | 0 | 0 | 1 | 0 | 0 |
| NW_015145035.1: 1402626-1409728 | 0.0107 | 0.0059 | -0.0047 | 9 | 0 | 0 | 5 | 0 | 0 |
| NW_015145135.1: 107689-121256 | 0.0071 | 0.0026 | -0.0045 | 11 | 0 | 0 | 4 | 0 | 0 |

**Table S6 - continued**

| | | | | | | | | | |
|---|---|---|---|---|---|---|---|---|---|
| NW_015145002.1: 2027477-2029020 | 0.0063 | 0.0018 | -0.0045 | 7 | 0 | 0 | 2 | 0 | 0 |
| NW_015145087.1: 1078823-1086391 | 0.0089 | 0.0045 | -0.0045 | 4 | 0 | 0 | 2 | 0 | 0 |
| NW_015145158.1: 751036-752074 | 0.0156 | 0.0111 | -0.0044 | 7 | 0 | 0 | 5 | 0 | 0 |
| NW_015145339.1: 214600-218862 | 0.0071 | 0.0027 | -0.0044 | 8 | 0 | 0 | 3 | 0 | 0 |
| NW_015145331.1: 656131-658106 | 0.0088 | 0.0044 | -0.0044 | 2 | 0 | 0 | 1 | 0 | 0 |
| NW_015145035.1: 1093225-1096999 | 0.0130 | 0.0087 | -0.0043 | 3 | 0 | 0 | 2 | 0 | 0 |
| NW_015145363.1: 420170-439338 | 0.0076 | 0.0032 | -0.0043 | 7 | 0 | 0 | 3 | 0 | 0 |

**Table S6 - continued**

| | | | | | | | | | |
|---|---|---|---|---|---|---|---|---|---|
| NW_015145088.1: 511895-515401 | 0.0086 | 0.0043 | -0.0043 | 4 | 0 | 0 | 2 | 0 | 0 |
| NW_015145036.1: 104626-110177 | 0.0129 | 0.0086 | -0.0043 | 6 | 0 | 0 | 4 | 0 | 0 |
| NW_015145003.1: 188898-190637 | 0.0107 | 0.0064 | -0.0043 | 5 | 0 | 0 | 3 | 0 | 0 |
| NW_015145002.1: 6038104-6039756 | 0.0043 | 0.0000 | -0.0043 | 1 | 0 | 0 | 0 | 0 | 0 |
| NW_015145039.1: 93280-93803 | 0.0043 | 0.0000 | -0.0043 | 1 | 0 | 0 | 0 | 0 | 0 |
| NW_015145400.1: 365022-369380 | 0.0085 | 0.0043 | -0.0043 | 8 | 0 | 0 | 4 | 0 | 0 |
| NW_015145002.1: 3263186-3267156 | 0.0170 | 0.0127 | -0.0042 | 16 | 0 | 0 | 12 | 0 | 0 |

**Table S6 - continued**

| | | | | | | | | | |
|---|---|---|---|---|---|---|---|---|---|
| NW_015145005.1: 5717202-5719747 | 0.0053 | 0.0011 | -0.0042 | 5 | 0 | 0 | 1 | 0 | 0 |
| NW_015145229.1: 768339-769157 | 0.0042 | 0.0000 | -0.0042 | 1 | 0 | 0 | 0 | 0 | 0 |
| NW_015145020.1: 1418226-1429916 | 0.0084 | 0.0042 | -0.0042 | 8 | 0 | 0 | 4 | 0 | 0 |
| NW_015145099.1: 1957413-1960415 | 0.0147 | 0.0105 | -0.0042 | 7 | 0 | 0 | 5 | 0 | 0 |
| NW_015145573.1: 101889-103914 | 0.0042 | 0.0000 | -0.0042 | 2 | 0 | 0 | 0 | 0 | 0 |
| NW_015145037.1: 1362652-1400957 | 0.0098 | 0.0056 | -0.0042 | 7 | 0 | 0 | 4 | 0 | 0 |
| NW_015145020.1: 2184591-2188447 | 0.0073 | 0.0031 | -0.0042 | 7 | 0 | 0 | 3 | 0 | 0 |

**Table S6 - continued**

| | | | | | | | | | |
|---|---|---|---|---|---|---|---|---|---|
| NW_015145028.1: 1454123-1459682 | 0.0063 | 0.0021 | -0.0042 | 3 | 0 | 0 | 1 | 0 | 0 |
| NW_015145339.1: 881829-884959 | 0.0083 | 0.0042 | -0.0042 | 6 | 0 | 0 | 3 | 0 | 0 |
| **Total** | | | | 259 | 0 | 0 | 111 | 0 | 2 |

FP = False Positive, FN = False Negative. Red highlighting corresponds to a tandemly-duplicated gene that was removed from the analysis.

**Table S7. SNP counts in *Diachasma* sequencing datasets.**

| wasp sample | Total SNPs | Homozygous SNPs | Heterozygous SNPs |
|---|---|---|---|
| D. muliebre I | 4013 | 3969 (98.90%) | 44 (1.10%) |
| D. muliebre II | 4005 | 3973 (99.20%) | 32 (0.80%) |
| D. muliebre III | 4013 | 3960 (98.68%) | 53 (1.32%) |
| D. muliebre IV | 4008 | 3964 (98.90%) | 44 (1.10%) |
| D. muliebre V | 3902 | 3863 (99.00%) | 39 (1.00%) |
| D. ferrugineum RNA | 3641 | 2888 (79.32%) | 753 (20.68%) |
| D. ferrugineum RRL | 3472 | 2830 (81.51%) | 642 (18.49%) |

**Total length of regions in overlap = 734,668 bp**

**Table S8. Statistical assessment of normality for genome datasets.**

| Wasp sample | Data type | # genes | W-statistic | p-value |
|---|---|---|---|---|
| D. ferrugineum | Pairwise distance | 3127 | 0.9650 | <2.2E-16 |
| D. muliebre I | Pairwise distance | 3127 | 0.9648 | <2.2E-16 |
| D. ferrugineum | Pairwise distance | 2978* | 0.9454 | <2.2E-16 |
| D. muliebre I | Pairwise distance | 2995* | 0.9467 | <2.2E-16 |
| D. ferrugineum | ENC | 3127 | 0.9389 | <2.2E-16 |
| D. muliebre I | ENC | 3127 | 0.9390 | <2.2E-16 |
| D. ferrugineum | CDC | 3127 | 0.9114 | <2.2E-16 |
| D. muliebre I | CDC | 3127 | 0.9115 | <2.2E-16 |
| D. ferrugineum | GC3 | 3127 | 0.9968 | 4.75E-6 |
| D. muliebre I | GC3 | 3127 | 0.9969 | 4.75E-6 |

ENC = effective number of codons, CDC = codon deviation coefficient, GC3 = GC content at third codon positions. *Gene counts with nonzero p-distance values.
